# The adipose-neural axis is critically involved in cardiac arrhythmias

**DOI:** 10.1101/2022.06.12.495845

**Authors:** Yubao Fan, Shanshan Huang, Suhua Li, Bingyuan Wu, Li Huang, Qi Zhao, Zhenda Zheng, Xujing Xie, Jia Liu, Weijun Huang, Jiaqi Sun, Xiulong Zhu, Maosheng Wang, Jieming Zhu, Andy Peng Xiang, Weiqiang Li

## Abstract

Dysfunction of the sympathetic nervous system and increase of epicardial adipose tissue (EAT) have been independently associated with the occurrence of cardiac arrhythmia. However, their exact roles in triggering arrhythmia remain elusive due to a lack of appropriate human disease models. Here, using the *in vitro* co-culture system with sympathetic neurons, cardiomyocytes, and adipocytes, we show that adipocyte-derived leptin could activate sympathetic neurons and increase the release of NPY, which in turn trigger arrhythmia of cardiomyocytes by interaction with NPY1R and subsequently enhancing the activity of NCX and CaMKII. The arrhythmic phenotype could be partially blocked by leptin neutralizing antibody, or an inhibitor of NPY1R, NCX or CaMKII. More importantly, increased EAT thickness accompanied with higher leptin/NPY blood levels was detected in atrial fibrillation patients compared to control group. Our study provides the first evidence that adipose-neural axis would contribute to arrhythmogenesis and represent a potential therapeutic target for arrhythmia.

## Introduction

Cardiac arrhythmia is one of the most common causes of morbidity and mortality and represents a major worldwide public health problem ^1^. Great progress has been made in managing different types of arrhythmias; the current therapies used to control heart rhythm include pharmacological therapies, catheter ablation, cardioversion/defibrillation, pacemaker, and autonomic therapies^1, 2^. However, some common and important rhythm disorders, such as atrial fibrillation (AF), ventricular tachycardia (VT), and ventricular fibrillation (VF), remain major therapeutic challenges because the underlying mechanisms are poorly understood ^1^. Critically, most of the current therapies are based on empirical rather than targeted mechanism-based approaches, and often lack sufficient efficacy and have considerable toxicity^3^. Therefore, more e□ective prevention and therapeutic options should be developed based on mechanistic data, with the goal of improving quality of life and/or reducing the symptom burden, hospitalization rate, and mortality in patients with arrhythmias.

Cardiac arrhythmias can result from structural or electrical abnormalities that lead to abnormal impulse formation and conduction, and are often associated with genetic or acquired heart diseases^4^. Previous studies reported that the sympathetic nervous system (SNS) plays an important role in the pathogenesis of cardiac arrhythmias^5^. Normally, the heart is densely innervated by SNS fibers, which regulate the rate, rhythm, and contractility of cardiomyocytes. However, aberrant SNS stimulation has been associated with AF, VT/VF, and sudden cardiac death through its ability to activate abnormal electrical pathways and increase the heterogeneity and dispersion of ventricular repolarization^6^. β-Adrenergic blockade is a cornerstone drug therapy for treating AF and VT/VF by targeting β-adrenergic receptors that respond to norepinephrine (NE). However, a significant proportion of patients continue to experience recurrent AF and VT/VF despite receiving the maximum tolerated dose of β-blockers^7^. This suggests that other, yet-unidentified stimulants are involved in cardiac arrhythmias, in addition to β-adrenergic agonists and the SNS.

In recent years, epicardial adipose tissue (EAT) thickness has been closely associated with the occurrence or recurrence of AF and VT/VF^5^. Under physiological conditions, EAT works as a buffer and functions as brown fat to supply energy and protect against hypothermia of myocardium^8^. EAT is in direct contact with the adjacent myocardium, without any barrier separating the tissues. Excess EAT may secrete various adipokines and inflammatory cytokines, which can cause inflammation and subsequent cardiac structural and electrical remodeling^9^. It has been hypothesized that pathological changes of EAT may be associated with abnormal autonomic tone augmentation and arrhythmias^9^. However, we lack direct evidence supporting the proarrhythmic properties of EAT, sympathetic neurons, and/or the interaction between them. This reflects a lack of appropriate human disease models and the difficulty of obtaining human cardiac tissue and/or propagating heart samples, sympathetic neurons, and primary adipocytes in culture^10^. Recently, stem cell technology has come to represent a powerful tool for developmental biology research, drug discovery, and the *in vitro* modeling of human disease. Cardiomyocytes generated from familial hypertrophic cardiomyopathy (HCM) patient-specific induced pluripotent stem cells *in vitro* were reported to display hypertrophy and electrophysiological irregularities, and provided clues that abnormal calcium handling properties may be involved in the development of HCM^11^. Sympathetic neurons and adipocytes were readily derived from human pluripotent stem cells (hPSCs) and adipose-derived stem cells (ADSCs), respectively^12, 13^. The above findings suggest that *in vitro* cardiac arrhythmia modeling could be performed using stem-cell derivatives, which may greatly facilitate the mechanistic study of cardiac disease pathogenesis.

In the present study, we generated sympathetic neurons, adipocytes, and cardiomyocytes from stem cells and established *in vitro* co-culture model systems to explore the effects of sympathetic neurons, adipocytes, or their interaction on cardiac rate and rhythm. We found that cardiomyocytes co-cultured with sympathetic neurons and adipocyte supernatant, but not either alone, exhibited a significant arrhythmic phenotype with electrical abnormalities and Ca^2+^ transient signaling irregularities. Our further experimental results suggested that adipocyte-secreted leptin may activate sympathetic neurons to enhance the release of NPY, which interacts with NPY Y1R on cardiomyocytes to eventually generate abnormal heart rhythms.

## Results

### Differentiation of functional sympathetic neurons from hPSCs

To generate the sympathetic neuron from stem cells *in vitro*, a stepwise induction protocol was established to transition hPSCs through the requisite intermediate stages of neural epithelial cells (NECs), neural crest stem cells (NCSCs), and sympathoadrenergic progenitors (SAPs; Fig. S1A). Monolayer cultured hPSCs treated for 5 days with N2B27 medium containing CHIR99021 (CHIR; activator of Wnt) and SB431542 (SB; inhibitor of GSK3β) were efficiently induced to differentiate into SOX1^+^/SOX2^+^/PAX6^+^/Nestin^+^ NECs (Fig. S1B). These day-5 (d5) NECs were cultured in suspension and treated with bone morphogenetic protein-2 (BMP2) and basic fibroblast growth factor (bFGF) to generate NCSCs. Fluorescence-activated cell sorting (FACS) analysis showed that the proportion of cells co-expressing the NCSC markers, p75 and HNK1, increased gradually during neural crest specification to exceed 95% on d11 (Fig. S1C). Most of the NCSCs highly co-expressed the trunk markers, HOXC8 and HOXC9, on d11 (Fig. S1D). We also observed substantial increases in the mRNA expression levels of HOX5-9 in differentiated cells treated with BMP2 and bFGF during neural crest commitment (from d7 to d11; Fig. S1E). It was previously reported that GD2 sorting can be used to enrich SAP-like cells derived from mouse and human ESCs^14^, and BMP signaling pathways have been implicated in the SNS-specific differentiation program^15^. Therefore, NCSCs were further incubated with medium containing BMP4. FACS analysis revealed that most of cells (94.35 ± 3.41%) co-expressed p75 and GD2 after 4 days of treatment with BMP4 (d15; Fig. S1F). We isolated GD2^+^/p75^+^ cells (SAPs) by FACS, and found that almost all of these cells co-expressed the SAP markers, PHOX2B, ASCL1, GATA3, HAND2, and PHOX2A (Fig. S1G, S1H).

We next explored whether SAP cells could be differentiated into sympathetic neurons using neural differentiation medium (Fig. 1A). After 4 weeks in culture, the cells showed a typical neuronal morphology (Fig. 1B), and large proportions of them stained positive for the catecholamine synthesis catalytic enzymes, tyrosine hydroxylase (TH, 48.20 ± 7.21%) and dopamine beta hydroxylase (DBH, 76.37 ± 5.44%). Additionally, over 40% of the differentiated cells were TH^+^/PHOX2B^+^ (45.47 ± 3.68%) or noradrenalin (NE)^+^/PHOX2B^+^ (40.1 ± 2.85%) (Fig. 1C). In accord with these immunocytochemical data, quantitative reverse transcription-polymerase chain reaction (qPCR) analysis showed that the mRNAs for the SAP lineage markers, ASCL1, PHOX2B, GATA3, TH, and DBH, were highly upregulated in SAP-like cells and sympathetic neurons (Fig. 1D). Fluo-4 acetoxymethyl ester (AM) (Fluo-4 AM) staining indicated that the *in vitro*-cultured sympathetic neurons exhibited spontaneous intracellular Ca^2+^ fluctuations and Ca^2+^ signaling could be evoked by the nicotinic acetylcholine receptor agonist, nicotine (Fig. 1E, F). Our whole-cell patch-clamp analysis showed that most of these neurons were capable of exhibiting spontaneous postsynaptic currents (82.50 ±1.53%, n=79) (Fig. 1G). Upon current injection, the neurons could fire a single action potential (AP) (Fig. 1H, left panel; Type □, 52.9 ± 1.73%, n=113) or a train of APs (Fig. 1H, right panel; Type □, 36.3 ± 1.14%, n=113) and exhibit a medium-size after-hyperpolarization (AHP) (Fig. 1I). We tested whether the depolarization of these hPSC-derived sympathetic neurons (SymNs) was dependent on Na^+^ ion channels and found that the APs of SymNs were completed blocked by 10 μM tetrodotoxin (TTX; a voltage-dependent Na^+^ channel blocker) (Fig. 1J). Taken together, these results demonstrate that functional sympathetic neurons could be efficiently generated using our protocol.

**Figure 1.**
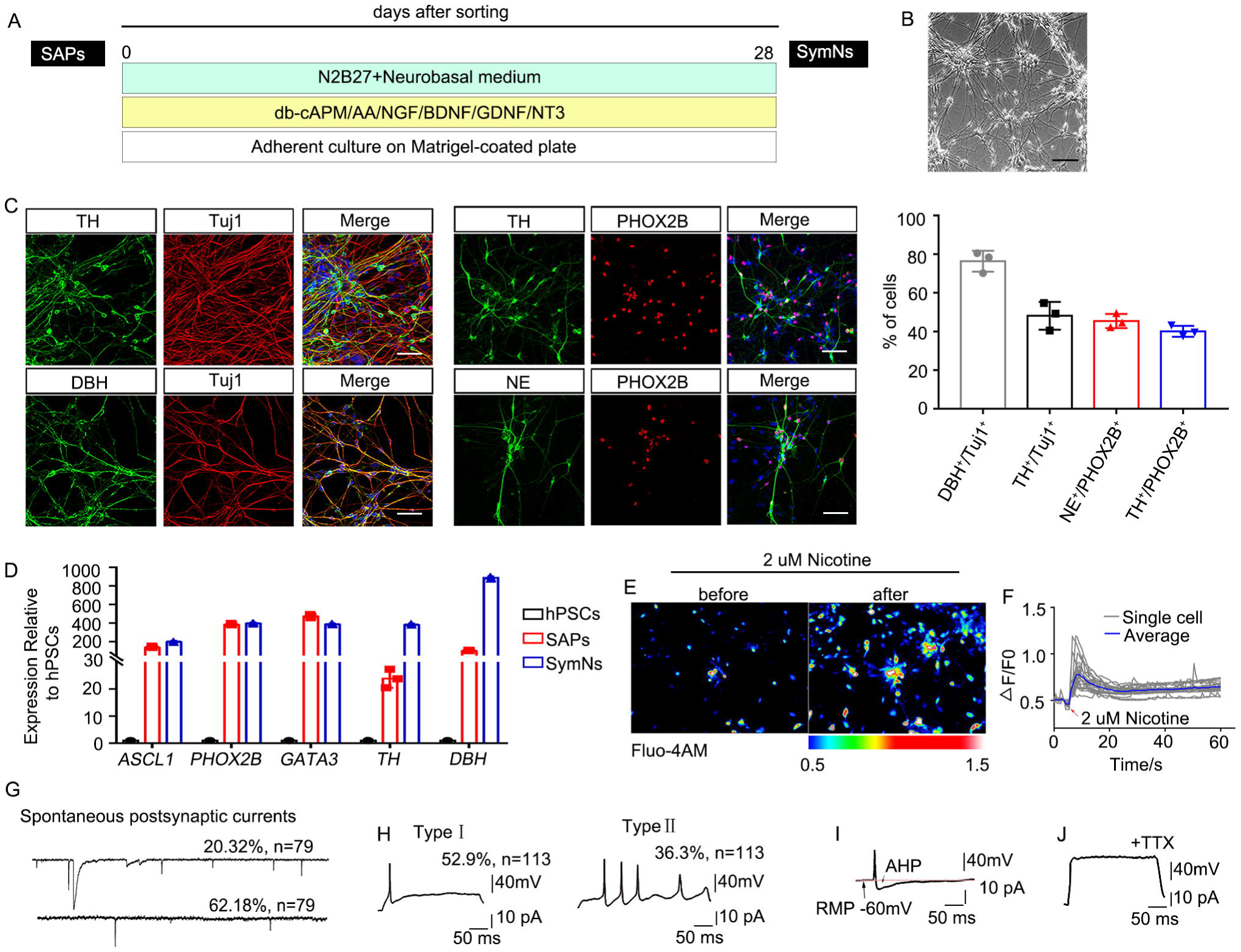
Characterization of sympathetic neurons derived from hPSC-SAPs. (A) Strategy for deriving sympathetic neurons from p75^+^/GD2^+^ hPSC-SAPs. (B) Representative bright field image of hPSC-derived sympathetic neurons. Scale bar, 50 μm. (C) The expression of sympathetic neuron markers including TH, DBH, Tuj1, NE, and PHOX2B in differentiated cells was detected by immunostaining, and the percentage of DBH^+^/Tuj1^+^, TH^+^/Tuj1^+^, NE^+^/PHOX2B^+^, and TH^+^/PHOX2B^+^ cells was quantified. Scale bar, 50 μm. (D) qPCR analysis for the mRNA expression of sympathetic neuron-specific genes during sympathetic neuronal differentiation from hPSCs. (E) The changes in intracellular calcium levels of sympathetic neurons induced by 2 μM nicotine were measured using Fluo-4AM. (F) Example traces of ΔF/F0 intensity of the sympathetic neurons exposed to 2 μM nicotine were recorded. Each trace is a response from a unique cell (n=22). (G) Spontaneous postsynaptic currents in sympathetic neurons were recorded with patch clamp techniques (62.18%, n=79). (H) A single (Type □; 52.9%, n=113) or a train (Type □; 36.3%, n=113) of action potential could be fired in sympathetic neurons. (I) An action potential evoked during a brief depolarizing current step, with an AHP. RMP, resting membrane potential. (J) The action potentials of sympathetic neurons were blocked by TTX. All of the error bars represent mean ± SEM.

### Cardiomyocytes exhibit an arrhythmia phenotype when co-cultured with SymNs and adipocyte supernatant

The sympathetic nervous system and EAT are thought to play important roles in cardiac electrophysiology and arrhythmogenesis^9^, but there is little direct evidence supporting this hypothesis. To investigate their exact roles in arrhythmia using in vitro cell model, we first derived cardiomyocytes from hPSCs using a commercial cardiomyocyte differentiation kit. Immunofluorescence and FACS analysis showed that more than 60% of the differentiated cell population represented cardiac troponin T (cTnT)-positive cardiomyocytes (73.28 ± 1.24%; Fig. S2A, B). We also induced adipose-derived mesenchymal stem cells (ADSCs) to differentiate to mature adipocytes (Fig. S2C). Our qPCR analyses showed that the ADSC-derived adipocytes expressed markers of white adipose tissue (WAT; UCP2, aP2, PPARγ, and C/EBPα), beige adipose tissue (BeAT; CD137, Cited1, Ear2, and TMEM26), and brown adipose tissue (BAT; UCP1, PRDM16, CPT1β, and C/EBPβ) (Fig. S2D). To determine whether EAT, sympathetic neurons, and/or their interaction might contribute to triggering arrhythmia, we established the following co-culture models: hPSC-derived cardiomyocytes (CMs) with hPSC-derived sympathetic neurons (SymNs) (CMs+SymNs), CMs co-cultured with ADSC-derived adipocyte supernatant (adi sup) (CMs+adi sup), and CMs co-cultured with SymNs and adi sup (CMs+SymNs+adi sup; triple co-culture system) (Fig. 2A-C). hPSC-CMs cultured alone were used as a control group. We found that the SymNs formed direct physical contacts with cardiomyocytes in the co-culture system (Fig. S3). Ca^2+^ was reported to play a fundamental role in regulating excitation-contraction coupling and electrophysiological signaling in the heart^16^. To investigate the rhythm of single cardiomyocytes, we used Fluo-4 AM to analyze the Ca^2+^ handling properties of control CMs and those in the different co-culture settings. Compared to control CMs, those co-cultured with SymNs showed a significantly faster beating rate (CMs: 15.50 ± 2.01 bpm, n=60; CMs+SymNs: 26.60 ± 2.80 bpm, n=60) and very few cells showing Ca^2+^ transient signal irregularity (CMs: 8.17 ± 2.64%, n=60; CMs+SymNs: 8.83 ± 2.48%, n=60) (Fig. 2D-F). Compared to control CMs, those co-cultured with adi sup showed similar beating rate (CMs: 16.33 ± 2.58 bpm, n=60; CMs+adi sup: 15.50 ± 1.87 bpm, n=60), and the percentage of Ca^2+^ transient irregularity increased slightly but not statistically significantly different from the control group (CMs: 8.17 ± 2.64%, n=60; CMs+adi sup: 14.00 ± 2.00%, n=60) (Fig. 2G-I). When co-cultured with both SymNs and adi sup, CMs exhibited a significantly higher beating rate (30.80 ± 4.11 bpm, n=60) and a strikingly large proportion of cells displaying Ca^2+^ transient irregularity (73.33 ± 6.25%, n=60); these irregularities included multiple events that may reflect arrhythmia-like voltage waveforms and were only rarely detected in control cells (Fig. 2J-L). However, CMs co-cultured with the supernatant from undifferentiated ADSCs showed a similar beating rate and percentage of cells showing Ca^2+^ transient irregularities compared to control CMs in the presence or absence of SymNs (data not shown). These findings indicate that the adipocyte supernatant may induce cardiomyocyte arrhythmia by activating the sympathetic neurons in our system.

**Figure 2.**
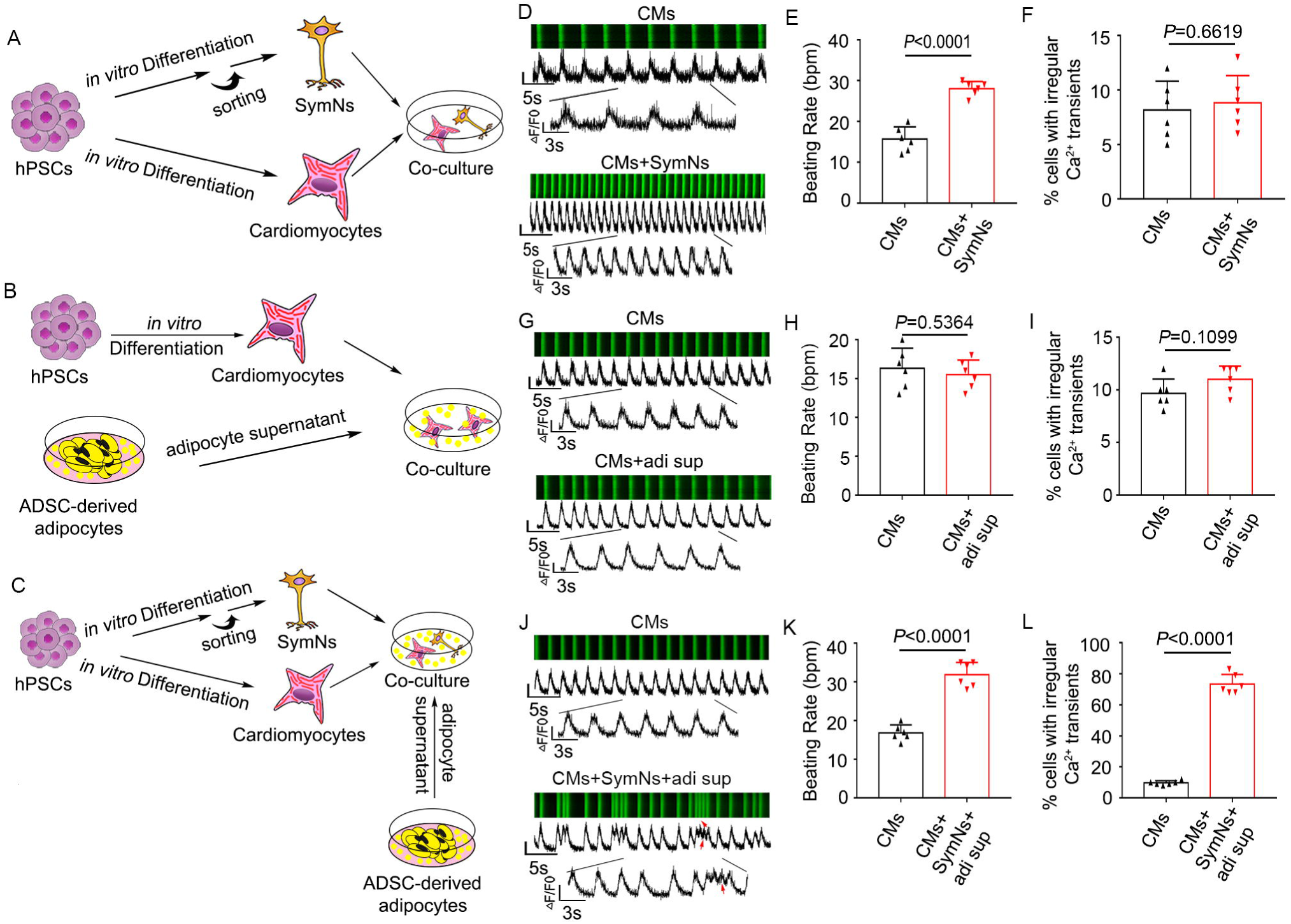
Adipocyte supernatant induces arrhythmic-like calcium transients in cardiomyocytes when cocultured with sympathetic neurons. (A) Schematic representation of the *in vitro* co-culture system using hPSC-derived cardiomyocytes and hPSC-derived sympathetic neurons (CMs+SymNs). (B) Schematic representation of the *in vitro* co-culture system using hPSC-derived cardiomyocytes co-culture and supernatant of ADSC-derived adipocyte (CMs+adi sup). (C) Schematic representation of the *in vitro* co-culture system using hPSC-derived cardiomyocytes, hPSC-derived sympathetic neurons, and supernatant of ADSC-derived adipocyte (CMs+SymNs+adi sup). (D) Representative line-scan images and spontaneous Ca^2+^ transients in CMs of control group and CMs+SymNs group. (E) Quantification of beating rate in CMs of control group and CMs+SymNs group. (F) Quantification of cells exhibiting irregular Ca^2+^ transients in CMs of control group and CMs+SymNs group. (G) Representative line-scan images and spontaneous Ca^2+^ transients in CMs of control group and CMs+adi sup group. (H) Quantification of beating rate in CMs of control group and CMs+adi sup group. (I) Quantification of cells exhibiting irregular Ca^2+^ transients in CMs of control group and CMs+adi sup group. (J) Representative line-scan images and spontaneous Ca^2+^ transients in CMs of control group and CMs+SymNs+adi sup group. Red arrows indicate arrhythmia-like waveforms. (K) Quantification of beating rate in CMs of control group and CMs+SymNs+adi sup group. Quantification of cells exhibiting irregular Ca^2+^ transients in CMs of control group and CMs+SymNs+adi sup group. All of the error bars represent mean ± SEM.

Next, we detected the field potential and conductivity of cardiomyocytes in our triple co-culture system by multiple microelectrode array (MEA) analysis (Fig. 3A, B). We found that the field potential duration (FPD) was significantly shorter for CMs cultured with SymNs compared with control CMs (CMs: 320.640 ± 17.412 ms; CMs+adi sup: 300.240 ± 16.540 ms; CMs+SymNs: 182.360 ± 6.973; CMs+SymNs+adi sup: 136.200 ± 12.157 ms), suggesting that the repolarization frequency and AF inducibility are increased in the CMs+SymNs and CMs+SymNs+adi sup groups (Fig. 3C, D). Similarly, the beat periods of single CMs clearly differed across the various groups, and the beat-to-beat variability of repolarization duration (BVR), which is negatively correlated with the autorhythmicity of cardiomyocytes, was sharply increased in the triple co-culture system (Fig. 3E). CMs of the triple co-culture system had multiple beat-starting points and displayed irregular electrical conduction compared with the other groups, indicating that CMs in this group not only had rhythm disorder at the single-cell level, but also exhibited the conduction of arrhythmia among CMs (Fig. 3F). These results imply that hPSC-CMs exhibited an arrhythmia phenotype in the CMs+SymNs+adi sup co-culture system.

**Figure 3.**
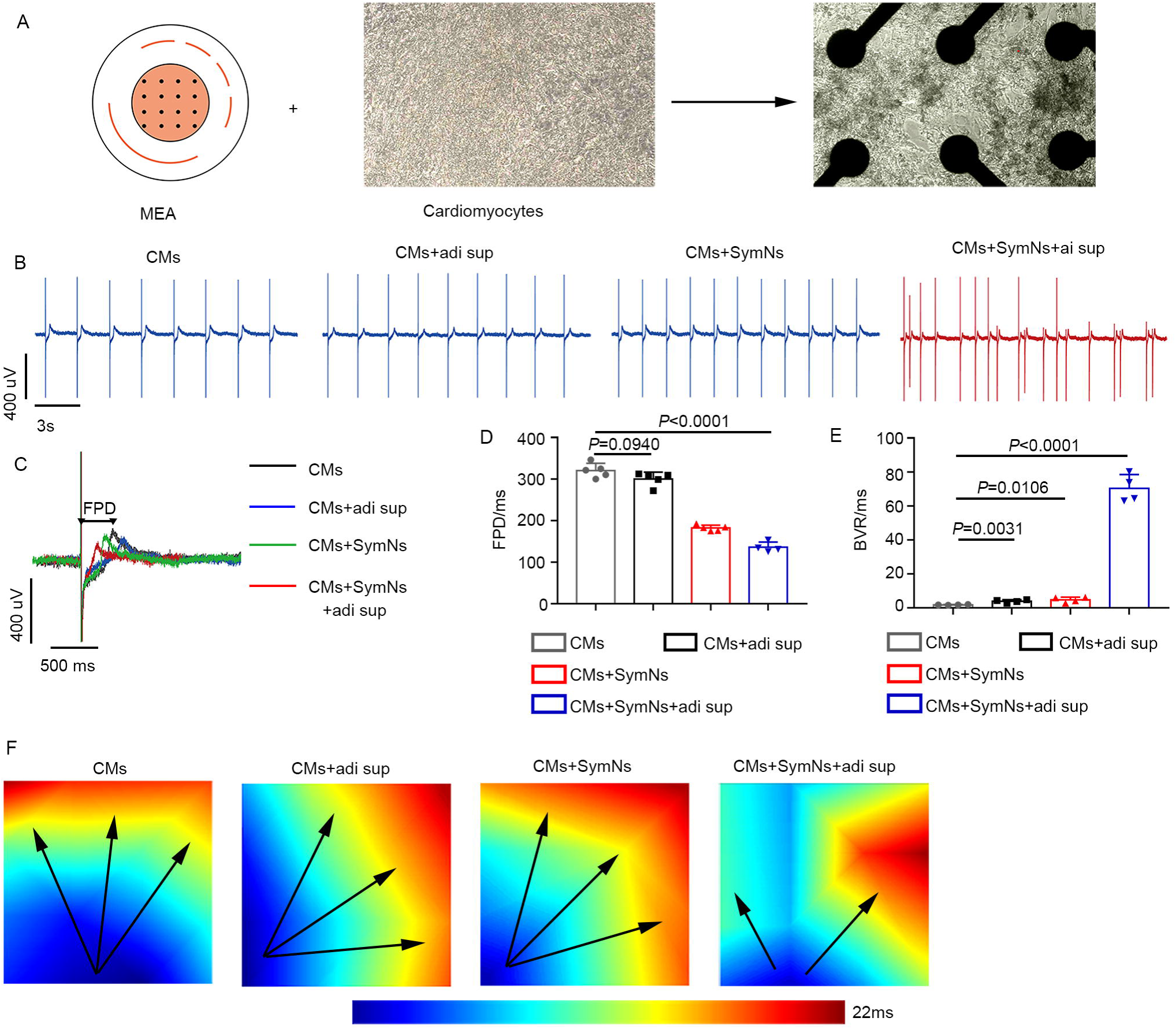
Detection of the arrhythmia phenotype of cardiomyocytes in triple co-culture system using MEA assay. (A) Schematic representation for MEA recordings of cardiomyocytes on an MEA plate. Cells are seeded by placing a droplet of the cell suspension directly onto the array and within the space circumscribed by the reference electrodes. (B) Representative traces of MEA recordings for cardiomyocytes in different groups. (C) Representative extracellular field potential recording of the cardiomyocyte in different groups. The arrows indicate the FPD. (D) Average values of FPD for each group were calculated and compared. (E) Average values of FPD BVR for each group were calculated and compared. (F) Typical conduction heatmap of cardiomyocytes in different groups. The blue region represents the origin of the beating (start electrode), while different colors showing the propagation delay time. The black arrows indicate the conduction direction of the filed action potential. All of the error bars represent mean ± SEM.

### Leptin secreted by adipocytes causes arrhythmia via activating sympathetic neurons

Adipocytes secrete various cytokines, including inflammatory cytokines and adipokines^17^. Here, we first used a bead-based multiplex assay to examine the amounts of different cytokines and adipokines in the supernatants of ADSCs (ADSC sup) or adipocytes differentiated from ADSCs (adi sup). Compared with undifferentiated ADSCs, the adipocytes expressed remarkably higher amounts of the inflammatory cytokines, IL-1β, IL-6, IL-8, and TNF-α, and the adipokines, adiponectin (APN) and leptin (Lep) (Fig. 4A). Adi sup also contained other cytokines (albeit at relatively lower levels), including IL-4, IL-10, CCL2, and TGF-β1 (Fig. S4). To test whether a single cytokine could alter the rhythm of cardiomyocytes by activating sympathetic neurons, we added each of the above-listed cytokines individually into the co-culture systems (CMs+cytokine or CMs+SymNs+cytokine) and then analyzed the Ca^2+^ transient signals of CMs using Fluo-4 AM. The beating rate of cardiomyocytes remained relatively constant under cytokine stimulation, regardless of the presence or absence of SymNs. Interestingly, the proportions of cells showing Ca^2+^ transient irregularities among CMs treated with a given single cytokine (10-15%) were comparable to that found in control CMs. However, the incidence of Ca^2+^ transient irregularities was considerably increased to about 20-25% in the CMs+SymNs+cytokine co-culture model (Fig. 4B-G). More strikingly, the addition of leptin (a proinflammatory adipocytokine secreted by adipocytes) resulted in extensive Ca^2+^ transient irregularities being seen in about 50% of CMs in the presence of SymNs (CMs+Lep: 15.50 ± 2.07%, n=60; CMs+SymNs+Lep: 51.50 ± 4.18%, n=60) (Fig. 4H, I). On the contrary, the addition of adiponectin, (a unique adipokine with multiple salutary effects such as antiapoptotic, anti-inflammatory, and anti-oxidative activities in numerous organs and cells^17^), did not significantly change the Ca^2+^ transient dynamics of CMs, regardless of the absence or presence of sympathetic neurons (Fig. 4J, K). Taken together, these results indicate that leptin may act through the adipose-neural axis to play an important role in the development of an arrhythmic phenotype.

**Figure 4.**
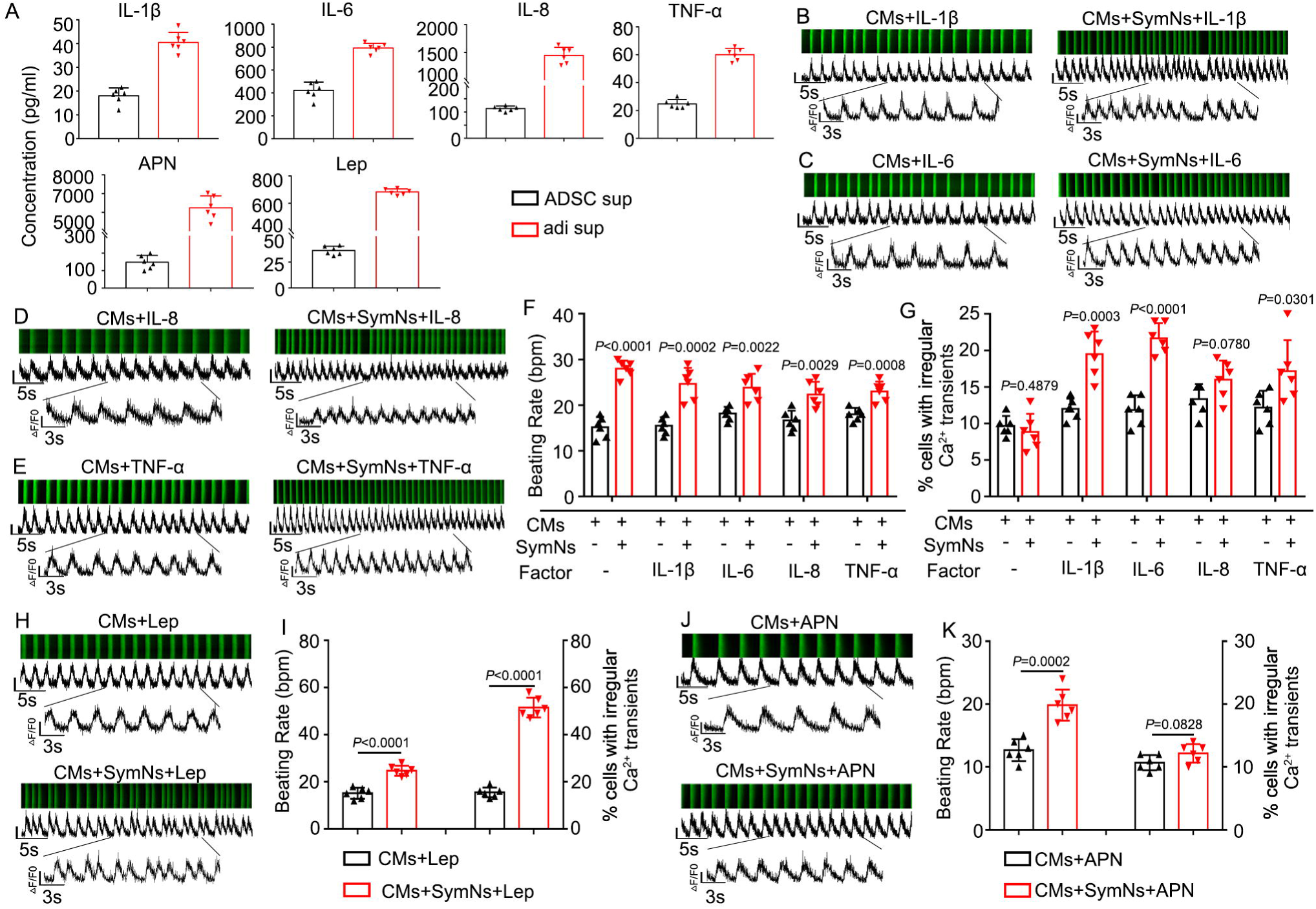
Leptin secreted by adipocytes causes arrhythmia in the presence of sympathetic neurons. (A) Quantification of cytokines in adipocyte supernatant using a bead-based multiplex assay. (B) Representative line-scan images and spontaneous Ca^2+^ transients in CMs of CMs+IL-1β group and CMs+SymNs+IL-1β group. (C) Representative line-scan images and spontaneous Ca^2+^ transients in CMs of CMs+IL-6 group and CMs+SymNs+IL-6 group. (D) Representative line-scan images and spontaneous Ca^2+^ transients in CMs of CMs+IL-8 group and CMs+SymNs+IL-8 group. (E) Representative line-scan images and spontaneous Ca^2+^ transients in CMs of CMs+TNF-α group and CMs+SymNs+TNF-α group. (F) Quantification of beating rate in CMs co-cultured with sympathetic neurons and a single cytokine. (G) The percentages of cells exhibiting irregular Ca^2+^ transients in CMs co-cultured with sympathetic neurons and a single cytokine. (H) Representative line-scan images and spontaneous Ca^2+^ transients in CMs of CMs+Lep group and CMs+SymNs+lep group. (I) Quantification of beating rate and percentages of cells exhibiting irregular Ca^2+^ transients in CMs of CMs+Lep group and CMs+SymNs+lep group. (J) Representative line-scan images and spontaneous Ca^2+^ transients in CMs of CMs+APN group and CMs+SymNs+APN group. (K) Quantification of beating rate and percentages of cells exhibiting irregular Ca^2+^ transients in CMs of CMs+APN group and CMs+SymNs+APN group. All of the error bars represent mean ± SEM.

We then examined the impact of leptin on sympathetic neurons. We found that leptin receptor (LepR) was abundantly expressed in sympathetic neurons (Fig. 5A), and treatment with leptin significantly increased the expression of c-Fos, a marker for neuronal activation, in TH-positive sympathetic neurons (Fig. 5B). Furthermore, ELISA revealed that the CMs+SymNs+Lep group released significantly higher levels of NE (CMs+SymNs: 201.00 ± 19.28 pg/ml; CMs+SymNs+Lep: 312.60 ± 13.29 pg/ml), NPY (CMs+SymNs: 388.30 ± 27.72 pg/ml; CMs+SymNs+Lep: 766.00 ± 10.31 pg/ml), and dopamine (DA; CMs+SymNs: 1.376 ± 0.06502 ng/ml; CMs+SymNs+Lep: 1.641 ± 0.05124 ng/ml) (Fig. 5C). Leptin was previously reported to activate the signal transduction molecule, STAT3, which in turn inhibits the expression of NPY in the brain areas responsible for controlling food intake. However, excessive or prolonged leptin treatment induces the expression of SOCS3, which is a negative feedback regulator of the STAT3 pathway^18^. Here, we found significantly downregulation of phosphorylated STAT3 and upregulation of SOCS3 (Fig. 5D) in SymNs after leptin treatment for 48 h. Similar results were obtained when hypothalamic neurons (mHypoA-1/2) were treated with leptin (Fig. S5A). To further confirm the role of leptin in the Ca^2+^ transient irregularities of CMs in our triple co-culture system, we used a neutralizing antibody to block the function of leptin present in the adipocyte supernatant. As expected, leptin neutralizing antibody treatment highly decreased the frequencies of Ca^2+^ transient irregularities in CMs (CMs+SymNs+Lep: 51.34 ± 5.13%, n=60; CMs+SymNs+adi sup + Lep nAbs: 28.83 ± 6.11%, n=60) (Fig. 5E, F). Moreover, the leptin-neutralizing antibody significantly reduced the levels of NE, NPY, and DA in the triple co-culture system (Fig. S5B). These findings suggest that cytokines secreted by adipocytes, especially leptin, could activate and increase neurotransmitter expression/release from SymNs via inhibition of the JAK2/STAT3 pathway.

**Figure 5.**
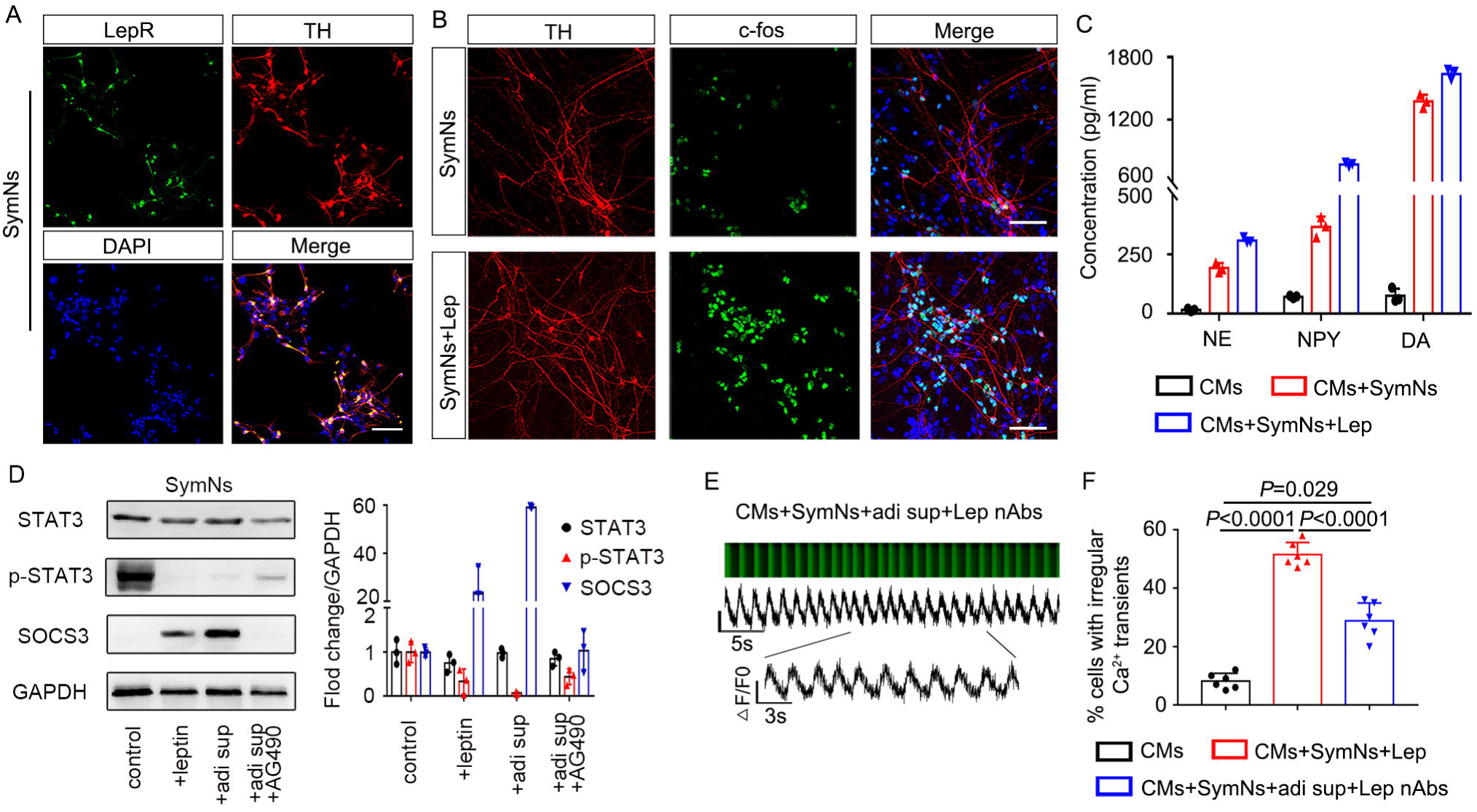
Leptin induces arrhythmia through activating the sympathetic neurons via leptin receptor. (A) Immunofluorescence staining for sympathetic neuron marker TH and the leptin receptor (LepR). Scale bar, 50 μm. (B) Immunofluorescence assay for TH and neuronal activity marker c-fos in sympathetic neurons cultured with or without leptin. Scale bar, 50 μm. (C) The concentrations of neurotransmitters in supernatant were analyzed using a commercial ELISA kit before and after leptin administration. (D) Western blot analysis and relative quantification for STAT3, p-STAT3, and SOCS3 in sympathetic neurons treated with leptin, adi sup, or adi sup plus JAK2 inhibitor (AG490). (E) Representative line-scan images and spontaneous Ca^2+^ transients in CMs of triple co-culture system with the addition of leptin neutralizing antibody. (F) Quantification of cells exhibiting irregular Ca^2+^ transients in CMs of triple co-culture system with the addition of leptin neutralizing antibody. All of the error bars represent mean ± SEM.

### The interaction of NPY and NPY1R causes arrhythmia by affecting NCX and CaMKII activity in the triple co-culture system

It has been reported that SNS could regulate cardiac function through adrenergic receptors expressed by cardiomyocytes^19^. Consistent with this, our qPCR results showed that hPSC-CMs highly expressed α-adrenergic receptors, β-adrenergic receptors, and dopaminergic receptors (Fig. S6A). The addition of phentolamine (α-adrenergic receptor inhibitor; 10 μM), propranolol (β-adrenergic receptor inhibitor; 10 μM), or L-741626 (dopamine D2 receptor inhibitor; 10 μM) was associated with little or no change in the frequency of Ca^2+^ transient irregularities in CMs co-cultured with sympathetic neurons and adipose supernatant (phentolamine: 77.17 ± 5.46%, n=60; propranolol: 59.50 ± 9.14%, n=60; L-741626: 68.167 ± 5.115, n=60) (Fig. 6A-F). These results suggest that the arrhythmia phenotype in CMs may not be mediated by NE/adrenergic receptor or DA/dopaminergic receptor interactions. Our qPCR results further showed that all five types of NPY receptor were expressed on cardiomyocytes, with Y1R, Y2R, and Y5R appearing at higher levels than Y3R or Y4R (Fig. 6G). The NPY mRNA was up-regulated in SymNs treated with leptin or adi sup (Fig. S6B), and the concentration of NPY in the supernatant increased time-dependently after SymNs were treated with leptin (Fig. S6C). Importantly, we found that addition of the Y1R antagonist, BIBP 3226 (1 μM), significantly reduced the frequency of Ca^2+^ transient irregularities in CMs co-cultured with SymNs and adi sup (22.17 ± 3.66%, n=60) (Fig. 6H-J), whereas the Y2R antagonist, BIIE 0246 (1 μM) did not substantially change the frequency of arrhythmic cardiomyocytes (67.50 ± 6.47%, n=60) (Fig. S6D-G). To further confirm the effect of NPY on cardiomyocyte arrhythmia, we treated cardiomyocytes with leptin, NPY, or Lep+NPY. Interestingly, we found that the groups treated with NPY and Lep+NPY had similar frequencies of CMs showing irregular Ca^2+^ transient signals (CMs+NPY: 66.500 ± 4.680, n=60; CMs+Lep+NPY: 64.835 ± 4.665, n=60), whereas leptin treatment alone did not increase the arrhythmia phenotype relative to that in the control CM group (CMs+Lep: 15.167 ± 2.858, n=60) (Fig. 6K, L). These results suggest that NPY, but not leptin, could directly induce an irregular rhythm among cardiomyocytes.

**Figure 6.**
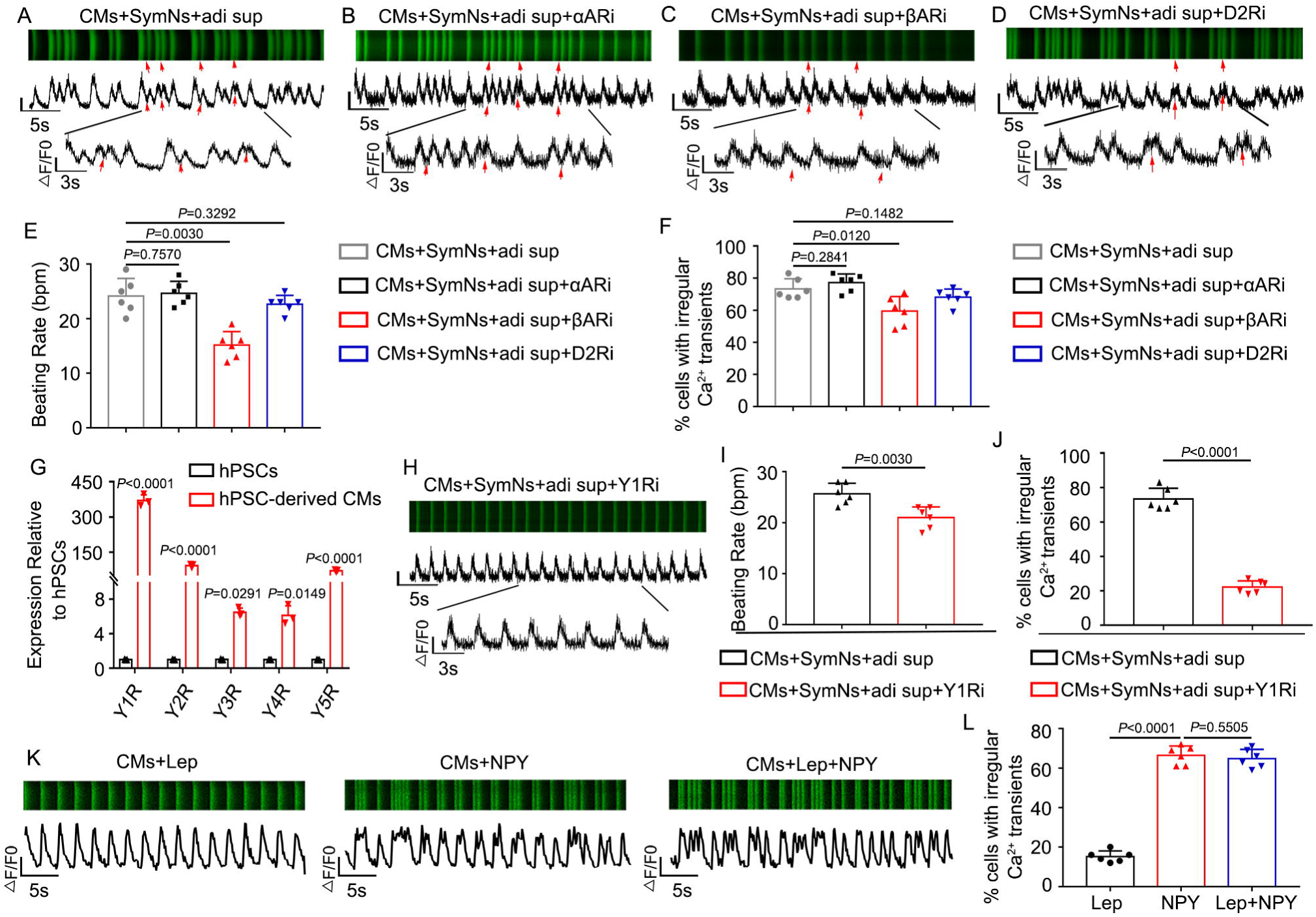
Activated sympathetic neurons lead to arrhythmia via NPY/Y1R interaction. (A) Representative line-scan images and spontaneous Ca^2+^ transients in CMs of triple co-culture system. Red arrows indicate arrhythmia-like waveforms. (B) Representative line-scan images and spontaneous Ca^2+^ transients in CMs of triple co-culture system with α-adrenergic receptors blocker, phentolamine. Red arrows indicate arrhythmia-like waveforms observed in this group similar to (A). (C) Representative line-scan images and spontaneous Ca^2+^ transients in CMs of triple co-culture system with β-adrenergic receptors blocker, propranolol. Red arrows indicate arrhythmia-like waveforms observed in this group similar to (A). (D) Representative line-scan images and spontaneous Ca^2+^ transients in CMs of triple co-culture system with D2R blocker, L-741626. Red arrows indicate arrhythmia-like waveforms observed in this group similar to (A) (E) Quantification of beating rate for CMs of triple co-culture system with a specific blocker. (F) Quantification of cells exhibiting irregular Ca^2+^ transients in CMs of triple co-culture system with a specific blocker. (G) qPCR analysis for the gene expression of NPY receptors in hPSC-derived cardiomyocytes. (H) Representative line-scan images and spontaneous Ca^2+^ transients in CMs of triple co-culture system with Y1R blocker (BIBP3226). (I) Quantification of beating rate in CMs of triple co-culture system with or without Y1R blocker. (J) Quantification of cells exhibiting irregular Ca^2+^ transients in CMs of triple co-culture system with Y1R blocker. (K) Representative line-scan images and spontaneous Ca^2+^ transients in CMs treatment with leptin, NPY or leptin plus NPY. (L) Quantification of cells exhibiting irregular Ca^2+^ transients in CMs treatment with leptin, NPY or leptin plus NPY. All of the error bars represent mean ± SEM.

One recent study reported that the NPY/Y1R interaction could induce arrhythmia^7, 20^, but the underlying mechanism remained unclear. Using a whole-cell patch clamp technique, we found that the action potential durations (APDs) at 30%, 50% and 90% repolarization levels (APD30, APD50, APD90) of the CMs+SymNs and CMs+SymNs+adi sup groups were shorter than that of the control group (for APD30, CMs: 299.9 ± 34.2 ms, n=9; CMs+SymNs: 101.7 ± 32.5 ms, n=9; CMs+SymNs+adi sup: 60.4 ± 10.8 ms, n=9; for APD50, CMs: 351.8 ± 33.5 ms, n=9; CMs+SymNs: 140.8 ± 42.2 ms, n=9; CMs+SymNs+adi sup: 82.2 ± 11.4 ms, n=9). This result indicated that CMs in the co-culture systems exhibited faster rhythms than control CMs, and was consistent with those observed for Ca^2+^ transient signaling. In contrast, APD30, APD50, and APD90 were increased in the CMs+SymNs+adi sup+Y1Ri group (APD30: 157.6 ± 36 ms, n=9; APD50: 209.6 ± 45.7 ms, n=9; APD90: 287.4 ± 39.8 ms, n=9) (Fig. 7A, B), suggesting that the Y1R inhibitor reduced outward currents and/or increased inward currents to slow the beating rate of CMs. Critically, analysis of AP traces revealed that the CMs+SymNs+adi sup group exhibited delayed after depolarizations (DADs; a typical arrhythmia feature) (Fig. 7C). The proportion of DADs occurring in the CMs+SymNs+adi sup group was 67.00 ± 5.65%; this decreased to 25.00 ± 4.24% in the presence of the Y1R inhibitor, indicating that the Y1R inhibitor reduced the occurrence of cardiac arrhythmia (Fig. 7D). DADs are often due to intracellular calcium overload, which enhances sodium-calcium exchanger (NCX) activity and promotes AF initiation^21^. Therefore, we designed a caffeine assay to identify whether NCX activity is enhanced in the triple co-culture system. Our results showed that the CMs+SymNs+adi sup group had the lowest calcium capacity and the strongest NCX activity, whereas these parameters were similar between the CMs+SymNs+adi sup+Y1R and CM groups (Fig. 7E-F).

**Figure 7.**
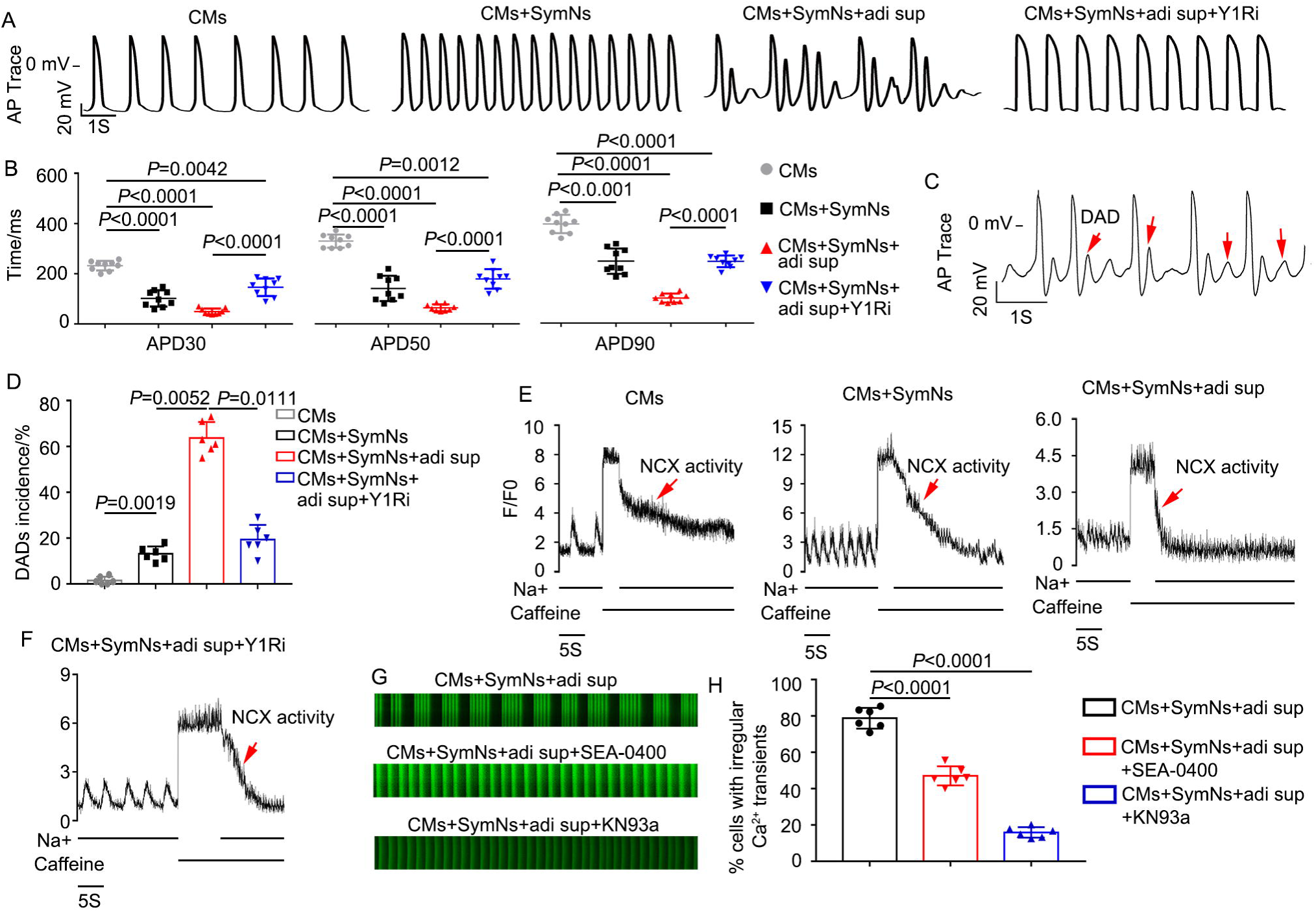
NPY/Y1R interaction results in arrhythmia through increasing NCX or CaMK□ activity. (A) Representative recordings of action potentials (APs) in CMs of indicated groups. (B) Average action potential durations (APD) at 30%, 50% and 90% repolarization levels were measured and compared (n=9). (C) Representative recordings of APs in CMs of triple co-culture system. Red arrows indicate typical DADs. (D) Quantification of DADs incidence for CMs in different groups. (E) Representative NCX-mediated calcium efflux in CMs of different groups during caffeine assay. The peak represents the thinness of SR Ca^2+^ loading. (F) Representative NCX-mediated calcium efflux in CMs of triple co-culture system with BIBP3226. The peak represents the thinness of SR Ca^2+^ loading. (G) Representative line-scan images and spontaneous Ca^2+^ transients in CMs of triple co-culture system with the NCX inhibitor (SEA0400; 2 μM, n=60) or the CaMK□ inhibitor (KN93a; 1 μM, n=60). (H) Quantification of cells exhibiting irregular Ca^2+^ transients in CMs of different groups. All of the error bars represent mean ± SEM.

The transient Ca^2+^ irregularity rate in the triple coculture system decreased from 78.77 ± 5.73% to 47.03 ± 5.24% upon treatment with the selective NCX inhibitor, SEA-0400 (Fig. 7G, H). Sustained activation of Ca^2+^/calmodulin-dependent protein kinase II (CaMKII) could also induce sustained DADs by increasing the spontaneous leakage of Ca^2+^ from the sarcoplasmic reticulum during diastole, which also plays a vital role in arrhythmia^22^. Our results revealed that the transient Ca^2+^ irregularity rate decreased prominently when KN93a (specific blocker of CaMK□) was added to the triple coculture model (CMs+SymNs+adi sup: 75.30 ± 5.38%; CMs+SymNs+adi sup+KN93: 14.17 ± 2.23%, n=60) (Fig. 7G-H). Taken together, these results indicate that the NPY/Y1R interaction plays a crucial role in adipose-neural axis-related arrhythmia by enhancing the activities of NCX and CaMKII.

### EAT thickness is positively correlated with the levels of leptin and NPY in coronary sinus blood from atrial fibrillation patients

To further investigate the correlation between human cardiac arrhythmia and EAT thickness or blood leptin/NPY levels, a total of 53 patients (58.7 ± 11.1 years; 64% male) were enrolled in this study, including 22 patients with paroxysmal AF, 17 with persistent AF, and 14 without AF (refer as control group). All patients were allowed to accept coronary computed tomography angiography (CCTA) to measure the EAT thickness, and underwent peripheral venous (PV) and coronary sinus (CS) blood sampling during electrophysiological diagnostic or therapeutic procedures. Among them, five AF patients further underwent open-heart surgery, and EAT samples were collected. The patients’ demographics, hemodynamics, and index of body fat are summarized in Supplementary Table 1 according to EAT thickness.

CCTA scan was used to measure the EAT thickness at different locations; we took the average EAT thickness at the left atrium (the average EAT thickness of periatrial-esophagus, periatrial-main pulmonary artery, and periatrial-thoracic descending aorta)^23^ for our statistical analyses (Fig. 8A). We also recorded the body mass index (BMI), subcutaneous adipose tissue (SAT) thickness, and abdominal fat (ABDF) thickness of recruited patients, and evaluated the correlation between EAT thickness and these indexes. Our results revealed that EAT thickness had weak positive correlations with BMI and SAT thickness (r<0.3) and weak or no correlation with ABDF thickness (r<0.2) (Fig. 8B, C; S7A). As expected, EAT thickness was significantly increased in patients with AF compared to those of the control group (Fig. 8D). Furthermore, we measured the leptin levels in patients and found that these levels were notably higher in CS blood compared to PV blood (Fig. 8E). We also found that the CS leptin level was more relevant to EAT thickness than the PV leptin level (Fig. 8F, G), and the CS leptin level was significantly higher in patients with AF compared to those of the control group (Fig. 8H). We then measured the NPY levels of PV or CS blood in patients. Although overall paired CS and PV NPY levels showed a significant positive correlation (r=0.9028), the concentration of NPY in CS blood was significantly higher than that in PV blood (Fig. 8I, J). Our results also showed that there were significant correlations between CS NPY and EAT thickness (r=0.8491) and between CS NPY and CS leptin (r=0.6842) (Fig. 8K, L). More importantly, higher NPY concentrations were detected in PV or CS blood of patients with AF compared to those of the control group (Fig. 8M; S7B). Finally, we tested the role of supernatants from AF patient-derived EAT (EAT sup) in our model. Consistent with the effects of adi sup, the presence of EAT sup was associated with a strikingly higher level of cardiomyocytes with transient Ca^2+^ irregularity in the triple co-culture system, compared with control CMs (CMs: 6.16 ± 1.46%; CMs+SymNs+EAT sup: 83.25 ± 3.49%, n=60). More importantly, this increase in the transient Ca^2+^ irregularity of cardiomyocytes could be partly mitigated by treatment with an Y1R inhibitor or leptin-neutralizing antibody (CMs+SymNs+EAT sup+Y1Ri: 18.55 ± 5.20%; CMs+SymNs+EAT sup+Lep nAbs: 41.46 ± 4.18%) (Fig. S7C, D). The above findings indicate that EAT thickness, but not SAT or ABDF thickness, is correlated with leptin and NPY in CS blood. Our results also suggest that, consistent with the results from our in vitro model, a high level of CS leptin and/or NPY might be an independent risk factor for AF patients.

**Figure 8.**
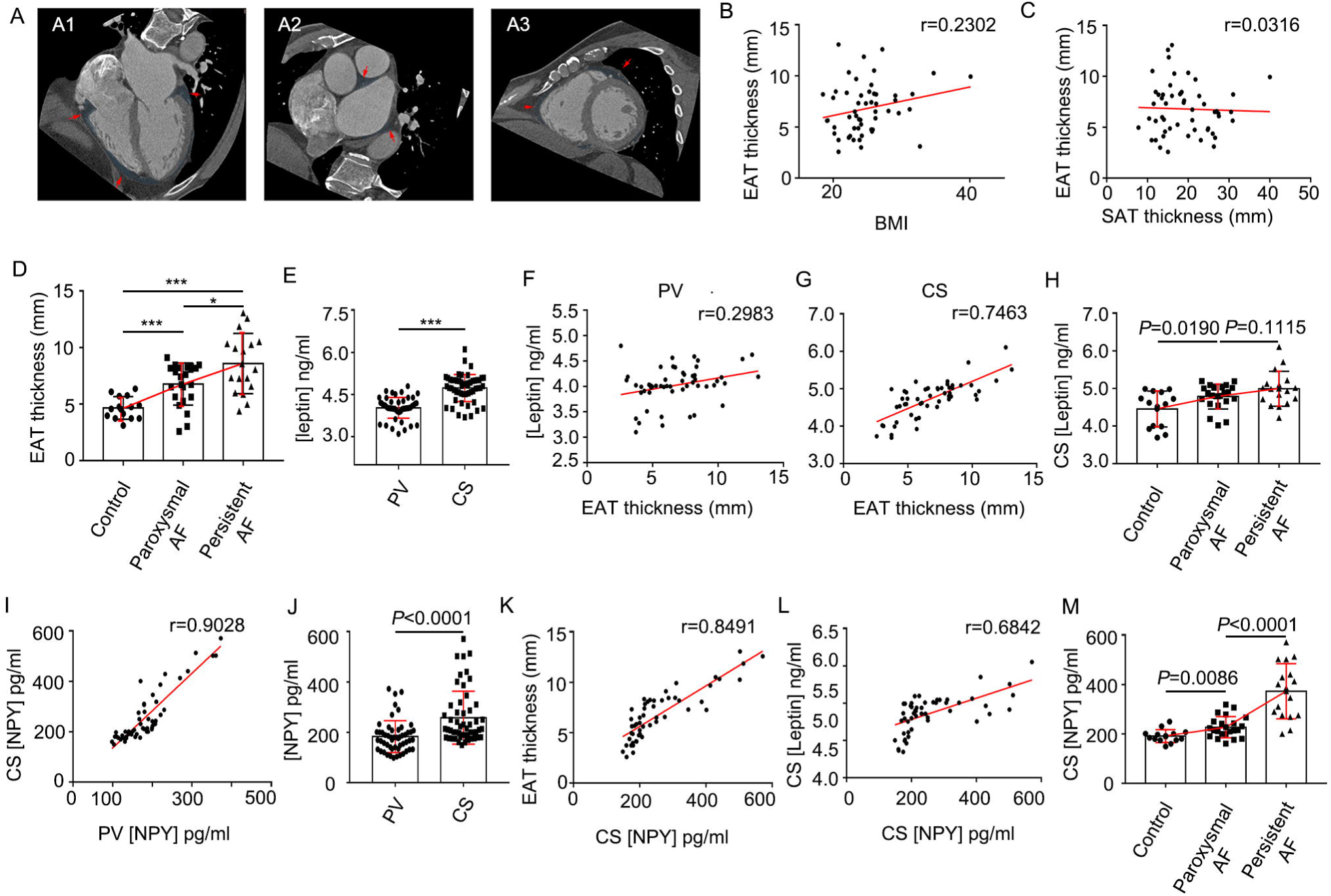
Arrhythmogenesis positively correlates with EAT thickness, leptin and NPY level of coronary sinus blood in human AF patients. (A) EAT thickness measurement by CT. A1: Horizontal long-axis plane for right atrioventricular groove (AVG), left AVG, and anterior interventricular groove (IVG); A2: Short-axis plane for periatrial EAT at esophagus, main pulmonary artery, and descending thoracic aorta; A3: Short-axis plane for superior IVG, inferior IVG, and right ventricular free wall. Red arrows indicate the point of measurement, and A2 showed the field of EAT that was used in this study. (B) Correlation between EAT thickness and BMI. (C) Correlation between EAT thickness and SAT thickness. (D) The EAT thickness were calculated and compared in patients with non-atrial fibrillation (control, n=14), paroxysmal atrial fibrillation (paroxysmal AF, n=22), and persistent atrial fibrillation (persistent AF, n=17). (E) Leptin concentrations in CS blood and PV blood were detected and compared. (F) Correlation between concentration of PV leptin and EAT thickness. (G) Correlation between concentration of CS leptin and EAT thickness. (H) The concentrations of coronary sinus leptin were detected and compared in patients with non-atrial fibrillation (control, n=14), paroxysmal atrial fibrillation (paroxysmal AF, n=22), or persistent atrial fibrillation (persistent AF, n=17). (I) NPY concentrations in CS blood and PV blood were detected and compared. (J) Correlation between PV- and CS-NPY levels across all patients. (K) Correlation between EAT thickness and concentration of CS NPY. (L) Correlation between concentration of CS leptin and CS NPY. (M) The concentrations of CS NPY were compared between AF patients with different severity. All of the error bars represent mean ± SD.

## Discussion

In the present study, we successfully established an in vitro co-culture system for arrhythmia modeling using hPSC-derived sympathetic neurons, cardiomyocytes, and ADSC-derived adipocytes. Our results revealed that when cardiomyocytes were co-cultured with sympathetic neurons and adipocyte supernatant, the frequency of cardiac arrhythmia increased strikingly. We also report for the first time that adipocyte-derived leptin could induce rhythm irregularity in cardiomyocytes by activating sympathetic neurons and promoting the release of NPY via enhancing the activities of NCX and CaMKII. Crucially, increased EAT thickness accompanied with higher levels of leptin and NPY were detected in AF patients compared to those of the control group, and supernatants from AF patient-derived EAT could act through leptin and NPY to induce Ca^2+^ transient irregularities in cardiomyocytes in the presence of sympathetic neurons.

Numerous studies revealed that sympathetic overactivation plays an important role in the pathophysiology of arrhythmias, and the abundance of EAT represents an independent risk factor for several types of arrhythmias^5^. Furthermore, it was speculated that EAT could contribute to triggering and maintaining arrhythmia through the modulation of autonomic activity^17^. However, most of these researches were correlation studies due to the lack of appropriate disease models. Here, we used cell co-culture system to simulate the in vivo cardiac microenvironment and investigated the critical role of adipose-neural axis in the pathogenesis of arrhythmias. We first showed that the beating rate of our CMs+SymNs co-culture model was much higher than that of the control, but the cardiac rhythm remained largely unchanged. These data indicate that the SNS might not induce cardiac arrhythmia before aberrant activation. Next, we found that adipocyte supernatant, AF patient-derived EAT supernatant, or a single relevant cytokine only caused a mild and not statistically significant increase in the occurrence of cardiomyocyte arrhythmias, indicating EAT alone also might not be the major initiator of arrhythmias. Intriguingly, when using the triple co-culture system (CMs+SymNs+adi sup), we found that the cardiomyocytes exhibited obvious rhythmic irregularities. We showed that adipocyte supernatant could induce the activation of sympathetic neurons and promote the secretion of major neurotransmitters. Our results further revealed that a single relevant proinflammatory cytokine existing in the adipocyte supernatant, especially leptin, could remarkably evoke rhythmic irregularity among cardiomyocytes in the presence of sympathetic neurons. These data provide the first evidence that the adipose-neural axis may contribute critically to arrhythmogenesis.

Leptin is an adipocyte-secreted hormone that regulates energy homeostasis, metabolism, neuroendocrine function, and other systems to suppress food intake and promote energy expenditure. Beyond its metabolic actions, leptin also stimulates the cardiovascular system by increasing sympathetic outflow via the renal, lumbar, and adrenal sympathetic nerves^24^. It is widely accepted that leptin modulates cardiac electrical properties and increases the heart rate by increasing sympathetic activity. In addition, leptin can directly decrease the heart rate, increase the QT interval, and induce ventricular arrhythmias in rats via leptin receptor, independent of β-adrenergic receptor stimulation^25^. In this study, we found that treatment with adipose supernatant or leptin alone did not change the rhythm of hPSC-CMs. These results were inconsistent with the previous reports, perhaps reflecting species-level differences. Our study further confirmed that leptin could induce arrhythmogenesis by directly activating sympathetic neurons and increasing the secretion of neurotransmitters, such as NE, DA, and NPY, in our triple co-culture system. Our results are consistent with a previous study showing that injection of leptin into the left stellate ganglion augmented ischemia-related ventricular arrhythmias via sympathetic nerve activation^26^. We found that the leptin level of CS blood was markedly higher in AF patients than in the control group, which further indicated that an interaction between local leptin in the heart and cardiac sympathetic neurons may play an important role in arrhythmogenesis. Furthermore, we demonstrated that the proarrhythmic role of adipocyte supernatant could be partially blocked by a leptin-neutralizing antibody, indicating there are other cytokines secreted by adipocytes may contribute to the arrhythmogenic properties by activating the SNS.

The sympathetic nervous co-transmitter, NPY, acts as an important regulator of cardiac function by directly interacting with NPY receptors on cardiomyocytes^7^. NPY can influence cardiovascular system function by acutely causing vasoconstriction, reducing acetylcholine release from the cardiac vagus, limiting subsequent bradycardia^27^, promoting ventricular and vascular remodeling, and participating in the pathogenesis of atherosclerosis^28^. More recently, the elegant study demonstrated NPY increased the incidence of ventricular arrhythmias in ST-elevation myocardial infarction patients, even in the presence of β-blockade^29^. Another study showed that NPY could reduce the threshold of occurrence for VT and VF^7^. In accordance with the above studies, we found that the NPY level was significantly increased in the triple co-culture system and in CS blood of AF patients, potentially accounting for the higher frequency of arrhythmic cardiomyocytes in the triple co-culture system compared to the control group. To further test whether NPY was responsible for the irregular cardiac rhythm observed in the triple co-culture system, we tested blockers for the NE, DA, and NPY signaling pathways. Our results showed that blockade of Y1R, but not α-adrenergic receptors, β-adrenergic receptors, dopamine D2 receptor, or Y2R, could efficiently rescue the cardiac arrhythmia seen in our triple co-culture system. These results supported the idea that NPY is pro-arrhythmic.

Delayed afterdepolarizations (DADs) are often due to calcium handling abnormalities caused by increased NCX or CaMK□ activity, and could induce the triggered activity at the cellular level and promote AF initiation^30^. To evaluate the cytosolic Ca^2+^ levels in CMs of the triple co-culture system, we analyzed the fluorescence intensity of calcium (proportional to the concentration of Ca^2+^) in the cytoplasm of CMs during the diastole and systole phases. Our preliminary data showed that during the diastole period, the calcium fluorescence intensity of the myocardium was significantly higher in the triple co-culture group than in control CMs, suggesting that co-culture system had a higher cytosolic calcium concentration and increased calcium leakage from the sarcoplasmic reticulum. We observed the opposite (decreased calcium fluorescence intensity of CMs in triple co-culture group compared with control CMs) during the systole stage. We also observed that the Ca^2+^ concentration tended to be down-regulated in the diastolic period and up-regulated in the systolic period in the presence of an Y1R blocker in the triple co-culture system (Fig. S8). Consistent with the above results, we found that NCX activity was upregulated in the presence of arrhythmia in the triple co-culture system, and this could be blocked by the Y1R inhibitor. The arrhythmia phenotype could also be partly suppressed by the NCX inhibitor. We further found that treatment with the specific CaMK□ blocker, KN93a, could significantly reduce the incidence of arrhythmia in the triple co-culture system. These data suggest that the interaction between NPY and Y1R induces an arrhythmic phenotype of cardiomyocytes by affecting NCX and CaMK □activity.

Taken together, these findings show that the adipose-neural axis plays a critical role in cardiac arrhythmia. We also demonstrate that the developed *in vitro* triple co-culture system can mimic the *in vivo* cardiac microenvironment and may supply a powerful tool for studying the pathogenesis of and exploring new prevention/treatment strategies for EAT/SNS-related diseases.

## Materials and methods

### Cell culture

Human pluripotent stem cells (hPSCs), including human induced pluripotent stem cells (hiPSCs) derived from human embryonic fibroblasts and human embryonic stem cells (hESCs; H9 cell line)^31, 32^, were used in this study. Undifferentiated hPSCs were cultured on Matrigel (Corning Inc., Corning, NY, USA)-coated culture plates with mTeSR1 medium (Stemcell Technologies, Vancouver, BC, Canada) as reported previously^33^. The medium was changed every day, and cells were passaged every 4-5 days using 0.5 mM EDTA (pH 8.0) (Thermo Fisher Scientific, Waltham, MA, USA).

### Neural crest and sympathoadrenal lineage induction from hPSCs

hPSCs were cultured to 80-90% confluence and harvested using Cell Dissociation Reagent (Accutase; Thermo Fisher Scientific) for 4-6 min at 37 □. For neural commitment, cells were seeded at 20,000-25,000 per cm^2^ onto Matrigel-coated culture dishes with mTeSR Plus complete medium supplemented with 10 μM Y-27632 (ROCK inhibitor; Sigma-Aldrich, St. Louis, MO, USA) and cultured at 37 □, 5% CO_2_ in a humidified incubator for 24h (day 0). On day 1, the medium was changed to N2B27 medium^34^ with 3 μM CHIR99021 (CHIR) and 10 μM SB431542 (SB) (both from Sigma-Aldrich) and cultured for 5 days. Medium was replaced on the day 3 after CHIR-SB induction. On day 5, the resultant cells were termed neural epithelial cells (NECs).

For neural crest differentiation, NECs were harvested using EDTA-PBS for 5 min and resuspended in Neurobasal medium (Thermo Fisher Scientific) consisting of 1% N2 (Thermo Fisher Scientific), 1% L-glutamine (1mM; Thermo Fisher Scientific), 10 ng/ml basic fibroblast growth factor (bFGF; Peprotech, Rocky Hill, New Jersey, USA), 10 ng/ml bone morphogenetic protein-2 (BMP2; Peprotech), 1% penicillin-streptomycin solution (Thermo Fisher Scientific) and 10 μM Y27632. Cells were cultured in suspension to form spheres in ultra-low attachment dishes for 6 days to generated neural crest stem cells (NCSCs; termed NCSCs-6d), and the medium was changed every other day.

For sympathoadrenergic progenitor (SAP) cell differentiation, the spheres of NCSCs-6d were harvested and re-plated onto dishes coated with Matrigel, and maintained in Neurobasal Medium consisting of 1% N2, 1% L-glutamine, 50 ng/ml bone morphogenetic protein-4 (BMP4; Peprotech), 1% penicillin-streptomycin solution (Thermo Fisher Scientific) for 4 days. Then cells were dissociated and labeled with antibodies against ganglioside GD2 (BD Pharmingen; Palo Alto, CA, USA) and low-affinity nerve growth factor receptor (NGFR, also known as p75; BD Pharmingen) for fluorescence-activated cell sorting (FACS). GD2 and p75 double-positive NCSCs were recognized as SAPs-like cells^35^. Immunocytochemistry and quantitative reverse transcription-polymerase chain reaction (qPCR) were used to analyze the expression of stem cell markers in differentiated cells.

### Sympathetic neuron differentiation from SAPs

For differentiation towards sympathetic neurons, GD2/p75 double-positive cells (SAPs) were plated onto glass coverslips in 48-well plates and cultured in differentiation medium containing DMEM-F12/Neurobasal medium (1:1 ratio; Thermo Fisher Scientific), 1% N2, 2% B27 (Thermo Fisher Scientific), 10 ng/ml brain-derived neurotrophic factor (BDNF; Peprotech), 10 ng/ml glial cell line-derived neurotrophic factor (GDNF; Peprotech), 20 ng/ml nerve growth factor (NGF; Peprotech), 20 ng/ml neurotrophin-3 (NT3; Peprotech), 200 μM ascorbic acid (AA; Sigma-Aldrich), and 0.5 mM dibutyryl-cAMP (db-cAMP; Sigma-Aldrich). Cells were differentiated for more than 4 weeks, and the medium was changed every 2-3 days. Differentiated cells were analyzed for the expression of sympathetic neural markers by immunocytochemistry and qPCR.

### Adipogenic differentiation of ADSCs

The ADSCs were purchased from American Type Culture Collection (ATCC, Washington, USA). For adipocyte differentiation, confluent ADSCs were treated with adipocyte differentiation medium (ADM) consisting of DMEM/High Glucose, 10% FBS (Thermo Fisher Scientific), 500 μM 3-isobutyl-1-methylxanthine (IBMX, Sigma-Aldrich), 2 μM Rosiglitazone (ROSI; Sigma-Aldrich), 1μM dexamethasone (Dex; Sigma-Aldrich), 10 μg/ml insulin (Sigma-Aldrich) for 6 days. The medium was changed every other day. After 6 days, the medium was replaced by ADM without IBMX and supplemented with Chemically Defined Lipid Concentrate (1:500; Thermo Fisher Scientific). Cells are maintained with daily medium change for additional 15-20 days. After differentiation, cells were analyzed by Oil Red O and qPCR. The adipocyte supernatant was collected and stored at −80□ for further study.

### Cardiac Differentiation of hPSCs

Cardiomyocytes (CMs) were generated from hPSCs was performed via WNT and retinoic acid (RA) signaling modulation. Briefly, about 3 × 10^5^ undifferentiated hPSCs were dissociated and re-plated onto a Matrigel-coated 12-well plate. Cells were first cultured with mTeSR Plus for 2 days (from day −2 to day 0) and expanded to 70% cell confluence. To initiate cell differentiation, the medium was changed to Cardiac Differentiation Medium (CDM; consisted of 1× Chemically Defined Lipid Concentrate, 10μg/ml Transferrin, 64mg/L L-ascorbic acid, and 13.6μg/L Sodium Selenium in DMEM/F12) supplemented with 5 μM CHIR99021 on day 1. The medium was changed to CDM with 0.6 U/ml Heparin (CDM+H) on day 2. From day 3 to day 5, the cells were treated with 3μM IWP2 and 1μM RA, and the medium was changed every day. From day 6 to day 8, the medium changed to CDM with Heparin again. The medium was changed to CDM with 20μg/ml insulin until beating was observed. Metabolic CM selection was performed using CDM without glucose, 0.5 mg/ml human recombinant albumin, and 4 mM lactate for two 3-days. hPSC-CMs of Day 30-40 after cardiac differentiation were utilized for downstream functional assays.

### Blood and epicardial adipose tissue sampling

Paired samples of coronary sinus (CS) and peripheral vein were collected from 53 patients after fully informed consent, including 22 patients with paroxysmal atrial fibrillation (AF), 17 patients with persistent AF, and 14 patients without AF (refer as control group). Cannulation of CS was performed via the femoral vein during electrophysiological diagnostic or therapeutic procedures. An 8F or 8.5F introducer sheath (St. Jude Medical, St. Paul, MN, USA) was placed into CS guided by a 6F steerable diagnostic catheter (St. Jude Medical). The position of the sheath was confirmed by CS angiography. Once the position was achieved, 10ml CS blood sample was drawn from the sheath. Another 10ml peripheral venous sample was collected at the same time. The samples were allowed to clot for 15 minutes and centrifuged immediately at 3300 rpm for 10 minutes. Samples were aliquoted and stored at −80°C until use. Leptin and NPY were measured using commercially available ELISAs (all from Millipore, Temecula, CA, USA) according to manufacturer’s instructions. Among the enrolled participants, five AF patients further underwent open-heart surgery, and epicardial adipose tissue (EAT) was collected. EAT was obtained following opening up of the pericardial sac with tissue collected over the body of the heart. Following tissue collection, the supernatant of EAT was collected and stored at −80℃ for further study. Ethical approval was obtained from the ethics committee of the Third Affiliated Hospital of Sun Yat-sen University (Approval No.: The Third Affiliated Hospital of Sun Yat-sen University [2020] 02-196-01).

### Ca^2+^ imaging of sympathetic neurons

Sympathetic neurons differentiated from hPSCs were used to evaluate the cytosolic Ca^2+^ signals. Coverslips containing sympathetic neurons were washed with warmed calcium buffer consisting of 130 mM NaCl (Sangon Biotech), 3 mM KCl (Sangon Biotech), 2.5 mM CaCl_2_ (Sangon Biotech), 0.6 mM MgCl_2_ (Sangon Biotech), 10 mM HEPES (Aladdin; Shanghai, CN), 10 mM Glucose (Sangon Biotech), 1.2 mM NaHCO_3_ (Sigma-Aldrich), adjusted to pH 7.2 with NaOH (Sangon Biotech). And then cells were incubated in warmed calcium buffer containing Calcium indicator, Fluo-4 acetoxymethylester (AM) (Thermo Fisher scientific) dissolved in DMSO (Sigma-Aldrich) (50 μg/μl) and 1% pluronic acid-127 (Sigma-Aldrich) for 15 min at 37℃. Cells were washed three times with warmed calcium buffer. Intracellular Fluo-4 AM was excited using automatic high-throughput living cell imaging and analysis system (BioTek, Lionheart FX, USA) at 494 nm. Baseline was monitored for 5 min and cells were stimulated with 2 μM nicotine (APExBio, Houston, TX, USA) to monitor for 10 min.

Alterations in time-dependent fluorescence were measured at a single wavelength. All analysis and processing, as well as playback of the image sequences for visual inspection, was made using ImageJ software (NIH, Bethesda, MD, USA). To visualize the changes in calcium signals resulting from spontaneous and stimulated activity, the raw sequences were processed to highlight changes in fluorescence intensity between frames and converted to heatmap, and the data was subsequently converted to a relative scale (ΔF/F baseline).

### Electrophysiology of sympathetic neurons

hPSC-derived sympathetic neurons cultured on coverslips were placed in a recording chamber on an inverted microscope and superfused with buffer containing 140 mM NaCl, 5 mM KCl, 10 mM glucose, 2 mM CaCl_2_ and 1 mM MgCl_2_ (all from Sangon Biotech), 10 mM HEPES (Aladdin), with pH adjusted to 7.4 using NaOH (Sangon Biotech), osmolarity of 300 mOsm with sucrose (Sangon Biotech). Electrodes were pulled from borosilicate glass (World Precision Instruments, Sarasota, FL, USA) and had resistance of 2-4 MΩ. The patch pipette was filled with intercellular solution containing 135 mM KCl, 1.1 mM CaCl_2,_ 10 mM HEPES (all from Sangon Biotech), 5 mM Na_2_-phosphocreatine, 2 mM EGTA, 2 mM Mg-ATP, 0.3 mM Na_2_GTP (all from Sigma-Aldrich), with pH adjusted to 7.3 using KOH (Sangon Biotech) and osmolarity adjusted to 290 mOsm using sucrose (Sangon Biotech). Tetrodotoxin (TTX; Sigma-Aldrich) was applied at a final concentration of 10 μM.

To assess parameters of neuronal excitability, resting membrane potential was monitored and cells were stimulated with 500-ms current injections ranging from −100 to +300 pA in 20-pA intervals, once every 5s. Steady-state inactivation was measured using 500 ms prepulses to voltages between −90 and 0 mV from a holding potential of −70 mV. Presynaptic APs were generated by passing current through a sharp recording electrode filled with 1 M K-acetate (70–90 MΩ), and EPSPs were recorded. Data were collected using Clampex 10.2 (Molecular Devices, Downtown, PA, USA).

### Ca^2+^ transient measurement of hPSC-CMs

hPSC-CMs were dissociated and seeded in Matrigel-coated Glass Bottom Cell Culture Dishes. Cells were loaded with 50 μg/μl Flou-4 AM and imaged in calcium buffer using a confocal microscope (LSM 710; Carl Zeiss, Jena, Germany). Spontaneous Ca^2+^ transients were recorded at 37℃ using standard line-scan methods. A total of 5000 lines scans were acquired for durations of 50 s. Average fluorescence intensity for Ca^2+^ line scans was quantified using Fiji (NIH). At least 60 cells in each group were recorded. The beating rates and the percentage of Ca^2+^ transient irregularities in each group were calculated by GraphPad prism 7.0 (CA, USA). NCX measurement was performed by flowing sodium free calcium buffer with 10 mM caffeine followed by calcium buffer with caffeine respectively. Cells were returned to normal calcium buffer at the end of recording. Timing between transients was defined as the time between the peaks of two successive spikes. The Ca^2+^ baseline was defined as the median of all minima of transients.

### Patch clamp recording of hPSC-CMs

To record cellular action potentials, hPSC-CMs were dissociated using STEMdiff Cardiomyocyte Dissociation Medium (Stemcell Technologies) for 15 min at 37℃, plated as single cells on 10 mm (48-well plate) glass cover slips coated with Matrigel in STEMdiff Cardiomyocyte Maintenance Medium and allowed to attach for 4-5 days. Cells were then subjected to whole-cell patch-clamp at 37℃, using an EPC-10 patch-clamp amplifier (HEKA) attached to a RC-26C recording chamber and mounted onto the stage of an inverted microscope (Nikon), and continuously perfused (2 mL/min) with Tyrode solution containing 140 mM NaCl, 4 mM KCl, 1.8 mM CaCl_2_, 1 mM MgCl_2_, 10 mM HEPES, 5.5 mM glucose (all from Sangon Biotech), with pH adjusted to 7.4 using NaOH. Patch-clamp experiments were performed in the whole-cell configuration. Electrodes were prepared as described above. The patch pipette was filled with internal solution containing: 30 mM KCl, 8 mM NaCl, 5 mM Mg-ATP, 10 mM HEPES, 0.01 mM EGTA (all from Sangon Biotech), with pH adjusted to 7.2 using KOH. Using this intracellular solution, the cells’ spontaneous beating, intracellular Ca^2+^ transient and contraction were preserved. Axopatch 200B amplifier (Axon Instruments Inc., Union City, CA, USA) was used for recordings.

### Multiplex assay

The cytokines in adipocytes supernatant were quantified by a bead-based multiplex assay (LEGENDplex^TM^ Human Panel; BioLegend, San Diego, CA, USA), according to the manufacturer’s instructions. Reactions were performed in three times. Eight key targets essential for immune response such as IL-1β, IL-6, IL-8, TNF-α, IL-4, IL-10, CCL2, TGF-β1, and adipokines including Adiponectin, and leptin were measured. Analysis was performed with a BD FACSCanto Ⅱ flow cytometer (BD Biosciences). The data was analyzed with LEGENDplex^TM^ V8.0 software (Biolegend).

### Multi-electrode array analysis (MEA)

Beating cardiomyocytes were re-plated in MEA plates (Axion Biosystems Inc., Atlanta, GA, USA) coated with 50 μg/ml fibronectin. When the cells generated rhythmic beating, the field potential (FP) waveform and contractibility were recorded for 10 min as baseline. Cells were co-cultured with sympathetic neurons and adipocytes supernatant for 2 days before the second MEA analysis. Field potential parameters including beating rate, filed potential duration (FPD) and conduction were analyzed and quantified using Cardiac analysis Tool (Axion Biosystems Inc.). 30 consecutive field potential periods were exported for beat-to-beat variability of repolarization duration (BVR) analysis in Clampfit 10.6 using the Axion Data Export Tool (Axion Biosystems Inc.) BVR was calculated using the following formula: 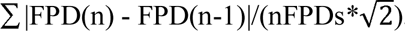.

### Western Blot analysis

Harvested cells were lysed in a Cell Total Protein Extraction Kit (KGP2100, KeyGen Biotech, Nanjing, China) and the protein concentration was measured by BCA assay. 20 μg of total protein was subjected to SDS-PAGE and transferred to PVDF membranes and then blocked for 1 hour with Tris-buffered saline containing 5% bovine serum albumin (BSA). Following blocking, the membranes were blotted for phospho-Stat3 (Cell Signaling Technology, Boston, USA, 1:1000), SOCS3 (Cell Signaling Technology, Boston, USA, 1:1000), and incubated at 4°C overnight, then washed and incubated with secondary antibodies at room temperature for 2 h. The signal was detected using the ChemiDoc Touch Imaging system (Bio-Rad, California, USA). All protein quantifications were normalized to the GAPDH expression level.

### Immunofluorescence analysis

Cells were fixed with 4% paraformaldehyde (PFA; Phygene, Gujing MaBao, FuZhou, CN) in phosphate-buffered saline (PBS) for 20 min. Then cells were treated with 0.3% Triton X-100 (Sangon Biotech; ShangHai, CN) for 40 min at room temperature and blocked with TBS containing 10% goat serum (Boster Biological Tech; CA, USA) or 5% donkey serum (Abbkine, Wuhan, CN.) for 1 hour at room temperature. Primary antibodies were incubated overnight at 4℃ followed by Alexa Fluor 488/594-conjugated goat anti-mouse/rabbit or Alexa Fluor 488-conjugated donkey anti-goat secondary antibodies for 2 hours at room temperature in the dark. Nuclei were counterstained with 4’,6-diamidino-2-phenylindole (DAPI; Sigma-Aldrich) for 5 min. Confocal images were obtained on Nikon C2 (Nikon; Nikon Eclipse Ni-E, JPN) laser scanning confocal microscope. All antibodies are listed in Supplemental Table 2.

### FACS analysis

Expression of cell surface markers in NCSCs and SAPs was determined by flow cytometry. Cells were harvested and incubated with monoclonal antibodies against human antigens, including p75, HNK1 and GD2 (all from BD Biosciences, San Jose, CA, USA). An irrelevant isotype-identical antibody (BD Biosciences) served as a negative control. Samples were analyzed by collecting 15,000 events using FlowJo software (BD Biosciences).

### Quantitative reverse transcription-polymerase chain reaction (qPCR) analyses

Total RNA was extracted using TRIzol Reagent (Thermo Fisher Scientific) from undifferentiated or differentiated cells. Genomic DNA was processed by DNase I (Fermentas, Glen Burnie, MD, USA) and complementary DNA synthesis was performed using Revert Aid First Strand cDNA Synthesis Kit (Thermo Fisher Scientific) using 1 μg of RNA. Gene expression was analyzed by LightCycler 480 SYBR Green Ⅰ Master kit with LightCycler 480 Detection System (Roche Diagnostics, Mannheim, Germany). qPCR values were normalized by GAPDH expression. The primer sequences used in this study are shown in Supplemental Table 3.

### ELISA

Cultured cells were washed twice with serum-free DMEM, and then the fresh medium was added. After 48h of incubation, the supernatant was gathered and centrifuged. The concentration of dopamine (DA), norepinephrine (NE) (both from abnova, Taipei City 114 Taiwan) and neuropeptide Y (NPY) (from Millipore) were detected by ELISA Kits (from Millipore) within 24h according to manufacturer instructions. The test of each sample was repeated at least three times.

### Statistical analysis

Demographic and clinical data are reported as the mean ± standard deviation (SD). Multiple linear regression analysis was used to determine the relationship between potentially inter-related variables. Experimental data were performed at least three times and showed as means ± standard error of mean (SEM). Comparisons between groups were performed using the paired or unpaired Student’s t-test or one-way analysis of variance (ANOVA). All statistical analyses were performed with the aid of SPSS Version 22 (SPSS Inc., Chicago, IL, USA). Graphs were drawn using GraphPad Prism 7.0.

## Acknowledgments

This work was supported by the National Key Research and Development Program of China (2018YFA0107200, 2021YFA1100603, 2019YFA0110303), the National Natural Science Foundation of China (32130046, 81970474, 81970222, 81901288), the Key Research and Development Program of Guangdong Province (2019B020234001, 2019B020236002).

## Conflict of interests

The authors declare that no competing interests exist.

## Contributions

W.L., A.P.X., and J.Z. conceived, organized and supervised the whole project. Y.F., S.H. performed cell acquisition and culture, established the co-culture model, collected and analyzed data. S.L., B.W., L.H., Z.Z., X.X., J.L., W.H., J.S., X.Z., and M.W. collected the clinical data and samples. Y.F. and B.W. measured the levels of NPY and Leptin in blood samples and analyzed the data. Y.F., S.H., and Q.Z. measured the electrophysiology of sympathetic neurons and cardiomyocytes. W.L., A.P.X., Y.F., and S.H. wrote the manuscript with the help from all the authors.

## Notes

### Competing Interest Statement

The authors have declared no competing interest.

