## Supplemental Table 1 for "The adipose-neural axis is critically involved in cardiac arrhythmias"

**Supplementary Table 1 Clinical characteristics according to EAT thickness**

|  | Normal EAT thickness (n=16) | | Increased EAT thickness (n=37) | | P-value |
| --- | --- | --- | --- | --- | --- |
| Age(years) | 57.6±3.3 | | 59.5±1.8 | 0.577 | |
| Males | | 6/16(37.5) | 28/37(75.7) | 0.0518 | |
| Cardiovascular risk factors | |  |  |  | |
| Hypertension | | 5/16(31.3) | 8/37(21.6) | 0.5314 | |
| Hyperlipidaemia | | 2/16(12.5) | 4/37(10.3) | 0.7742 | |
| Diabetes mellius | | 4/16(25.0) | 3/37(8.11) | 0.0056 | |
| Blood pressure and heart rate | |  |  |  | |
| Systolic(mmHg) | | 124.4±4.3 | 125.9±2.7 | 0.7537 | |
| Diastolic (mmHg) | | 73.8±2.0 | 81.1±1.6 | 0.01 | |
| .leart rate (/min) | | 69.6±4.1 | 80.5±4.0 | 0.1073 | |
| Type of AF | |  |  |  | |
| Non-AF | | 10/16(62.5) | 4/37(10.8) | 0.0015 | |
| Paroxysmal AF | | 4/16(25.0) | 18/37(48.6) | 0.0034 | |
| Persistent AF | | 2/16(12.5) | 15/37(40.5) | 0.0013 | |
| BMI | | 23.6±0.8 | 25.4±0.7 | 0.1682 | |
| SAT thickness (mm) | | 18.5±1.6 | 19.3±1.1 | 0.6842 | |
| ABDF thickness (mm) | | 12.0±1.0 | 12.6±0.8 | 0.6695 | |
| Values are expressed mean ± SD or n (%); AF, atrial fibillation. | | | |  | |
