## Supplemental Table 2 for "The adipose-neural axis is critically involved in cardiac arrhythmias"

**Supplementary Table 1 Primary antibodies used in immunostaining**

| Primary antibodies | Host | Dilution | Cat.No. | Supplier |
| --- | --- | --- | --- | --- |
| PAX6 | Rabbit | 1:350 | ab195045 | Abcam, Cambridge, UK |
| SOX2 | Rabbit | 1:500 | Ab97959 | Abcam, Cambridge, UK |
| SOX1 | Goat | 1:200 | AF3369 | R&D Systems, Minneapolis, MN, USA |
| Nestin | Rabbit | 1:200 | Mab353 | Millipore, Temecula, CA, USA |
| HOXC8 | Rabbit | 1:200 | ab79690 | Abcam, Cambridge, UK |
| HOXC9 | Mouse | 1:50 | ab50839 | Abcam, Cambridge, UK |
| PHOX2B | Mouse | 1:50 | SC-376997 | Santa Cruz,Delaware Ave,CA,USA |
| ASCL1 | Rabbit | 1:100 | ab211327 | Abcam, Cambridge, UK |
| HAND2 | Rabbit | 1:50 | ab200040 | Abcam, Cambridge, UK |
| GATA3 | Rabbit | 1:250 | ab199428 | Abcam, Cambridge, UK |
| PHOX2A | Mouse | 1:50 | Sc-81978 | Santa Cruz,Delaware Ave,CA,USA |
| TUBB3 | Mouse | 1:1000 | mab1195 | R&D Systems, Minneapolis, MN, USA |
| TH | Rabbit | 1:500 | ab152 | Millipore, Temecula, CA, USA |
| DBH | Rabbit | 1:1000 | Mab308 | Millipore, Temecula, CA, USAC |
| NE | Rabbit | 1:500 | ab8887 | Abcam, Cambridge, UK |
| c-Fos | Mouse | 1:100 | Ab208942 | Abcam, Cambridge, UK |
| LeptinR | Goat | 1:50 | AF389 | R&D Systems, Minneapolis, MN, USA |
| cTnT | Mouse | 1:100 | MA5-12960 | ThermoFisher Scientific, Waltham, MA,USA |
