## Supplemental Table 3 for "The adipose-neural axis is critically involved in cardiac arrhythmias"

| **Supplementary table 2 primers for qPCR** | | |
| --- | --- | --- |
| **Gene** | | **Primers** |
| GAPDH | Forward | GAAGGTGAAGGTCGGAGTC |
|  | Reverse | GAAGATGGTGATGGGATTTC |
| HOXB5 | Forward | AGCTCCAGCGCCAATTTCA |
|  | Reverse | TCCGGCCCGGTCATATCAT |
| HOXB7 | Forward | GACCCCGGCAATTTCTACGG |
|  | Reverse | ATCCAGGGGAAGAGCTGTGT |
| HOXC5 | Forward | CTAAGAGCAGTGGGGAGATCA |
|  | Reverse | GTCATCCACGGGTAAATCTGTG |
| HOXD9 | Forward | GGTTGGGGTTTGTCCTCAGT |
|  | Reverse | TTTACAACTGGTCCTCGGGC |
| PHOX2B | Forward | ACGCCGCAGTTCCTTACAAA |
|  | Reverse | CTGGTGAAAGTGGTGCGGAT |
| ASCL1 | Forward | TCCCCCAACTACTCCAACGA |
|  | Reverse | GCGATCACCCTGCTTCCAAA |
| GATA3 | Forward | TAACATCGACGGTCAAGGCAAC |
|  | Reverse | GTAGGGATCCATGAAGCAGAGG |
| TH | Forward | GTTCTCCCAGGACATTGGACTT |
|  | Reverse | ACACAGCCCAAACTCCACA |
| DBH | Forward | CCTGACCCTGCTTTTCAAGAG |
|  | Reverse | GTTCGGGGATATTGGGCTTCA |
| UCP2 | Forward | CCCCGAAGCCTCTACAATGG |
|  | Reverse | CTGAGCTTGGAATCGGACCTT |
| aP2 | Forward | ATGGGGGTGTCCTGGTACAT |
|  | Reverse | ACGCCTTTCATGACGCATTC |
| C/EBPα | Forward | TCGGTGGACAAGAACAGCAA |
|  | Reverse | TTGTCACTGGTCAGCTCCAG |
| PPARγ | Forward | TGAAGACGGATTGCCCTCATT |
|  | Reverse | GCTGGTGCCAGTAAGAGCTT |
| CD137 | Forward | TTGGATGGAAAGTCTGTGCTTG |
|  | Reverse | AGGAGATGATCTGCGGAGAGT |
| Cited1 | Forward | GCTGGCTAGTATGCACCTGC |
|  | Reverse | CATTGGCTCGGTCCAACCC |
| TMEM26 | Forward | GAAGATGGCTTCTACCCATTGG |
|  | Reverse | GTGCAGGACTATTCCTCACATTT |
| Ear2 | Forward | GGCCCTCACCGAGTATGTG |
|  | Reverse | CAGTGTCTCAATGGGCGTCT |
| UCP1 | Forward | TGTCCTGGGAACAATCACCG |
|  | Reverse | GTGCTGTTTCTTTCCCTGCG |
| C/EBPβ | Forward | AAGCACAGCGACGAGTACAA |
|  | Reverse | ACAGCTGCTCCACCTTCTTC |
| CPT1β | Forward | CTTTCTTCGTGGCCCTGGAT |
|  | Reverse | CGCATGCTCTGCATTGAGAC |
| PRDM16 | Forward | CTTCGGATGGGAGCAAATACTG |
|  | Reverse | TCCACGCAGAACTTCTCACTG |
| NPY | Forward | CGCTGCGACACTACATCAAC |
|  | Reverse | CTCTGGGCTGGATCGTTTTCC |
| ADRβ1 | Forward | ATCGAGACCCTGTGTGTCATT |
|  | Reverse | GTAGAAGGAGACTACGGACGAG |
| ADRβ2 | Forward | TGGTGTGGATTGTGTCAGGC |
|  | Reverse | GGCTTGGTTCGTGAAGAAGTC |
| ADRβ3 | Forward | GACCAACGTGTTCGTGACTTC |
|  | Reverse | GCACAGGGTTTCGATGCTG |
| Y1R | Forward | GAGGCGATGTGTAAGTTGAATCC |
|  | Reverse | TGGAACGGCTCATCAGTCATT |
| Y2R | Forward | CATCTTGCTTGGGGTAATTGGC |
|  | Reverse | AGAGTGAACGGTAGACACAGAG |
| Y3R | Forward | CGGAGCTGAGGAGAACGAC |
|  | Reverse | ATAGCCCGTGCCAATCACC |
| Y4R | Forward | GTGCCTTACAGCGTTTCAGAG |
|  | Reverse | ACCCTTGGCATATCTCAGTGA |
| Y5R | Forward | CGGTAAACTTCCTCATAGGCAAT |
|  | Reverse | AACATCCACTGATCCAGCAAG |
| DRD1 | Forward | CTCCGTTTCCAAATACATTCCA |
|  | Reverse | CACTGTTGATTCTTTGCCCT |
| DRD2 | Forward | AGCATCGACAGGTACACAG |
|  | Reverse | CTCGTTCTGGTCTGCGT |
| DRD3 | Forward | GTGGTATACCTGGAGGTGAC |
|  | Reverse | GCAGTGTACCTGTCTATGCT |
| DRD4 | Forward | CCGCTCTTCGTCTACTC |
|  | Reverse | ACAGGTTGAAGATGGAGG |
| DRD5 | Forward | CTCATCTCCTACAACCAAGAC |
|  | Reverse | TGATAGATCTGGAACATGCGA |
| ADRα1 | Forward | TCTGGCGGTCATTCTAGTCAT |
|  | Reverse | GGTGTCCTCGTGAAAGTTCTTG |
| ADRα2 | Forward | AGAGGTCAACGGACACTCGAA |
|  | Reverse | CCCCACAAACACCCTCCTT |
| ADRα3 | Forward | ACAAGAGAGAACCAGACTCCAA |
|  | Reverse | AGGGTAGTTAAACTCCTCCTCC |
