## supplementary figures for "The adipose-neural axis is critically involved in cardiac arrhythmias"

**Supplemental Figures and Figure legends**

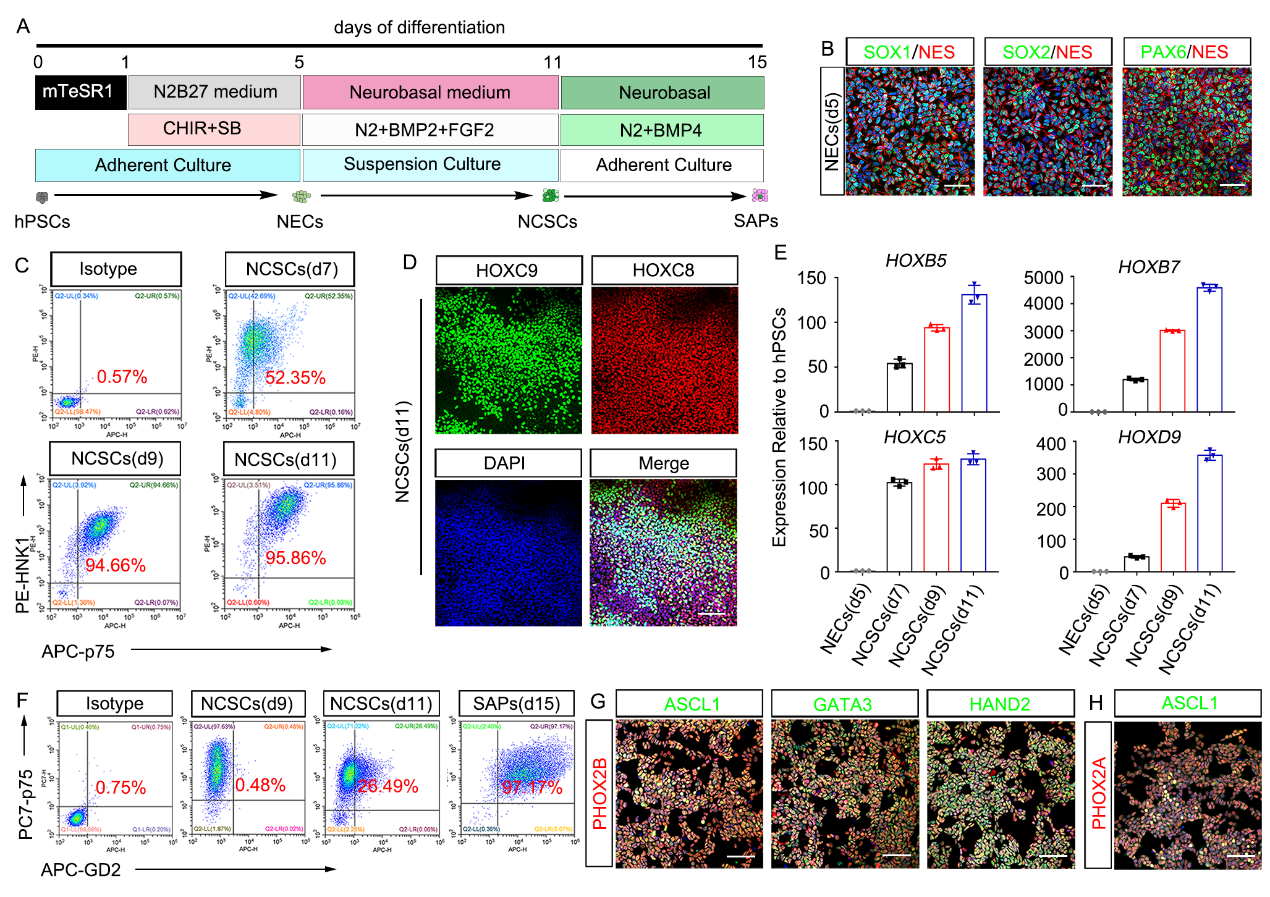

**Figure S1 In vitro differentiation of human pluripotent stem cells toward SAPs.**

(A) Schematic diagram for differentiation of hPSCs into SAPs.
(B) Immunofluorescence assay for the expression of NEC markers SOX1, SOX2, PAX6, and NES in d5 differentiated cells. Scale bar, 50 μm.
(C) FACS analysis for NCSC markers (p75 and HNK1) in differentiated cells on d7, d9, and d11.
(D) Immunofluorescence staining of trunk neural crest markers HOXC8 and HOXC9 in d11 differentiated cells. Scale bar, 50 μm.
(E) qPCR analysis for the expression of HOX genes in differentiated cells on d5, d7, d9, and d11.
(F) FACS analysis for SAP markers (p75 and GD2) in differentiated cells on d9, d11 and d15.
(G) Immunofluorescence staining of SAP markers ASCL1, PHOX2B, GATA3, and HADN2 in p75^+^/GD2^+^ SAPs. Scale bar, 50 μm.
(H) Immunofluorescence staining of SAP markers ASCL1 and PHOX2A in p75^+^/GD2^+^ SAPs. Scale bar, 50 μm.

All of the error bars represent mean ± SEM.

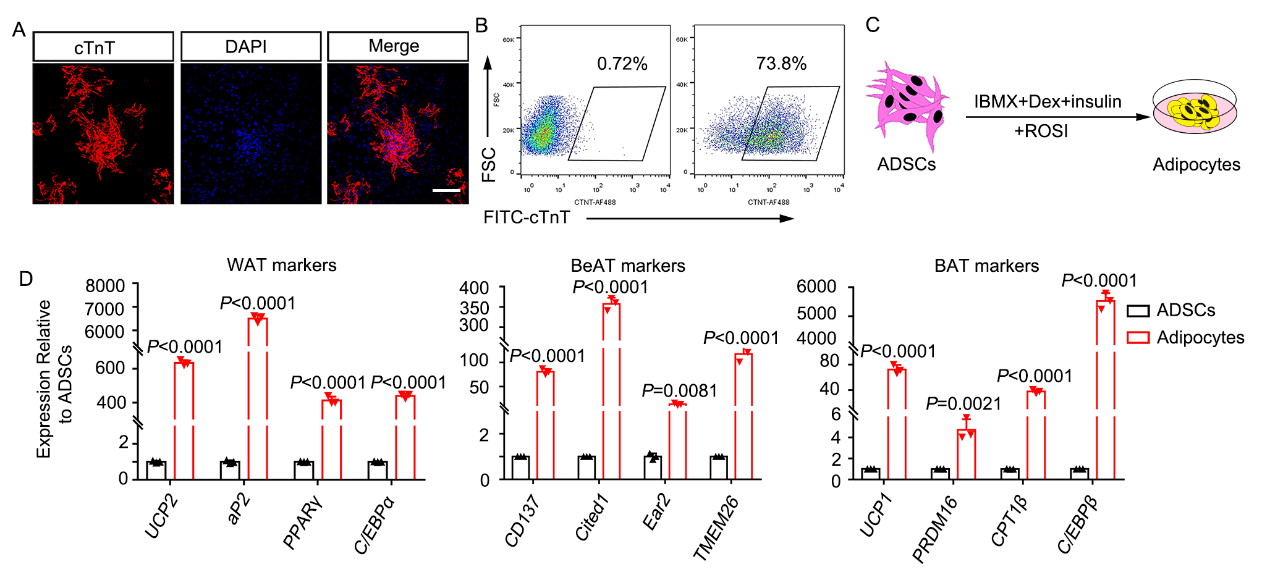

**Figure S2 Identification of hPSC-derived CMs and ADSC-derived adipocytes.**

(A) Immunofluorescence analysis for cTnT expression in hPSC-derived CMs. Scale bar, 50 μm.
(B) FACS analysis for the expression of cTnT in hPSC-derived CMs.
(C) Schematic diagram for adipocyte differentiation from ADSCs.
(D) qPCR analysis for WAT/BeAT/BAT markers expressed in undifferentiated ADSCs and ADSC-derived adipocytes.

All of the error bars represent mean ± SEM.

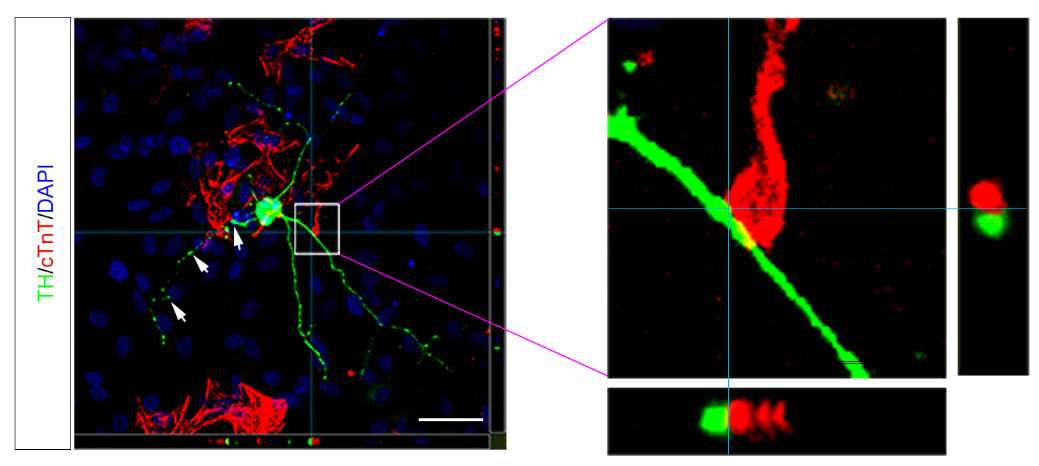

**Figure S3 The expression of sympathetic neurons marker TH and cardiomyocyte marker cTnT in co-culture system were detected by immunofluorescence assay.** Scale bar, 50 μm.

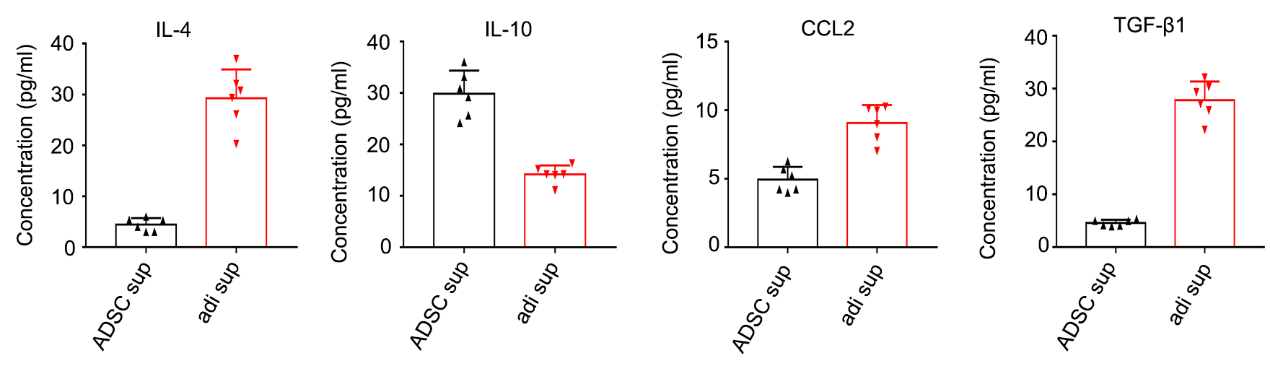

**Figure S4 Quantification of cytokines in adipocyte supernatant using a bead-based multiplex assay.**

All of the error bars represent mean ± SEM.

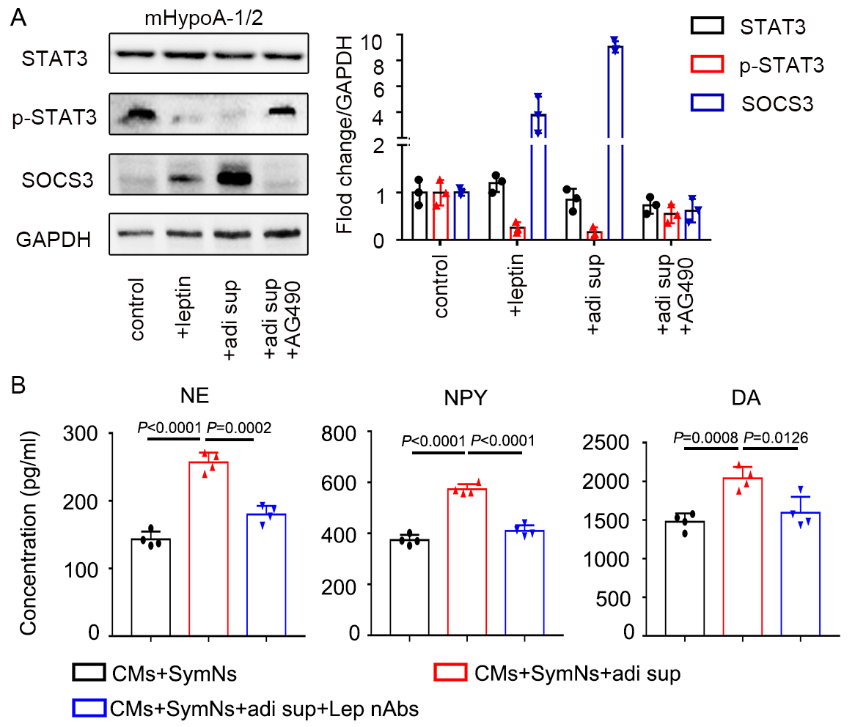

**Figure S5 Leptin activates hypothalamic neurons through JAK2-STAT3 pathway and leptin neutralizing antibody could block the activation of sympathetic neurons in triple co-culture system.**

(A) Western blot analysis and relative quantification for STAT3, p-STAT3 and SOCS3 in mHypoA-1/2 hypothalamic cell line treated with leptin, adi sup, or adi sup plus JAK2 inhibitor (AG490).
(B) The concentrations of neurotransmitters in supernatant were detected using a commercial ELISA kit before and after addition of leptin neutralizing antibody.

All of the error bars represent mean ± SEM.

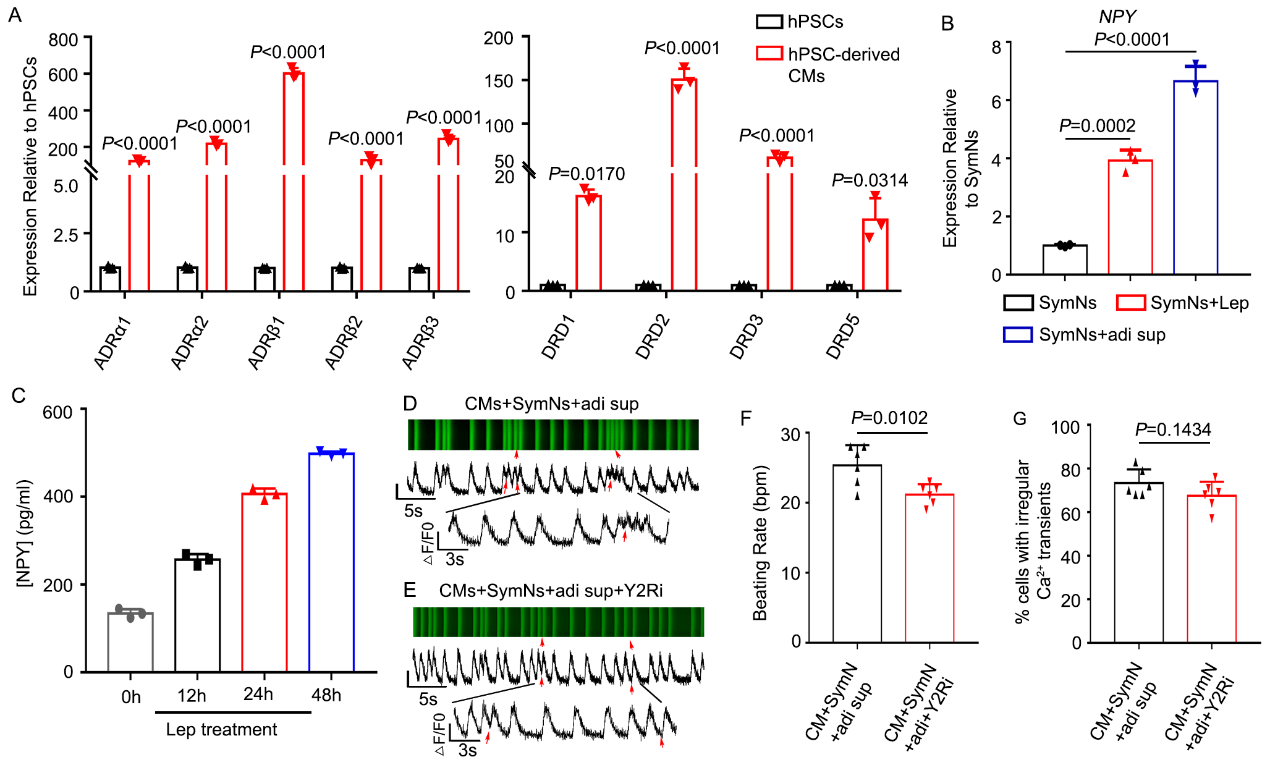

**Figure S6 Cardiomyocytes express adrenergic and dopaminergic receptors, and the arrhythmia of cardiomyocytes in triple co-culture system is not mediated by NPY/NPY2R interaction.**

(A) qPCR analysis for gene expressions of adrenergic and dopaminergic receptors in hPSC-derived cardiomyocytes.
(B) qPCR analysis for NPY mRNA expression in sympathetic neurons after treatment with leptin or adi sup.
(C) The concentration of NPY in supernatant after leptin treatment was analyzed at different time points by ELISA assay.
(D) Representative line-scan images and spontaneous Ca^2+^ transients in CMs of triple co-culture system. Red arrows indicate arrhythmia-like waveforms.
(E) Representative line-scan images and spontaneous Ca^2+^ transients in CMs of triple co-culture system with Y2R blocker (BIIE 0246). Red arrows indicate arrhythmia-like waveforms.
(F) Quantification of beating rate in CMs of triple co-culture system with Y2R blocker.
(G) Quantification of cells exhibiting irregular Ca^2+^ transients in CMs of triple co-culture system with Y2R blocker.

All of the error bars represent mean ± SEM.

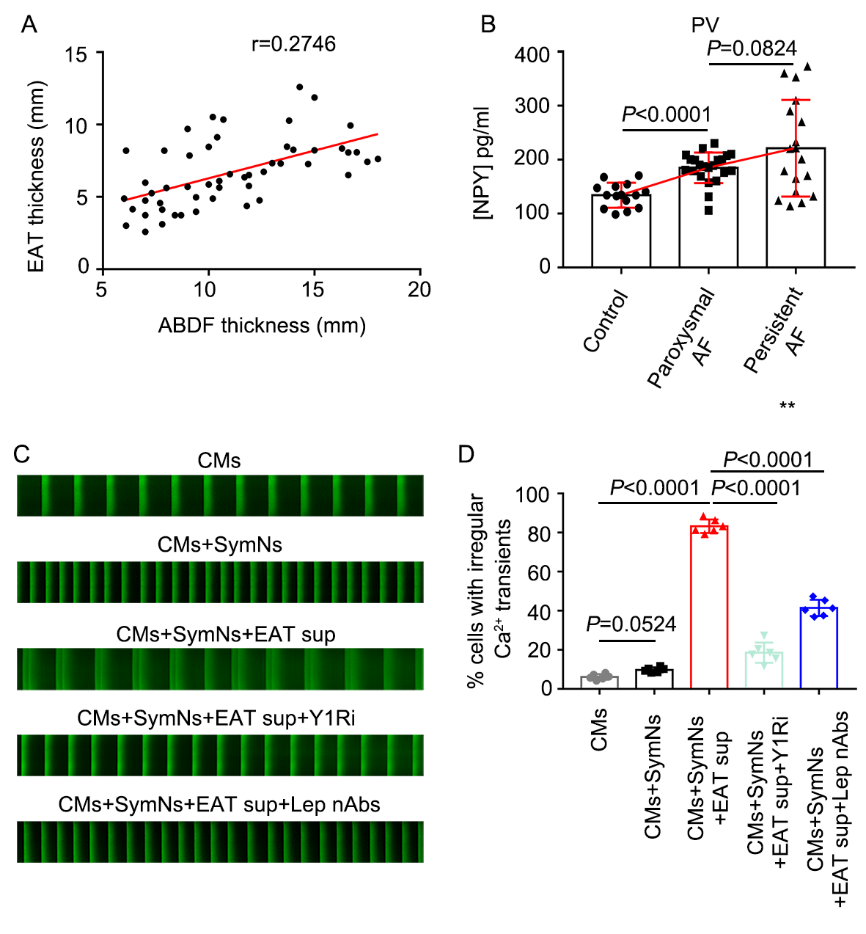

**Figure S7 Arrhythmogenesis positively correlates with NPY level in PV blood in human AF patients, and human EAT supernatant-related arrhythmia could be suppressed by leptin neutralizing antibody and Y1R blocker.**

(A) Correlation between EAT thickness and ABDF thickness.
(B) The concentrations of PV NPY were compared between AF patients with different severity.
(C) Representative line-scan images and spontaneous Ca^2+^ transients in CMs of different groups.
(D) Quantification of cells exhibiting irregular Ca^2+^ transients in CMs of different groups.

All of the error bars represent mean ± SEM.

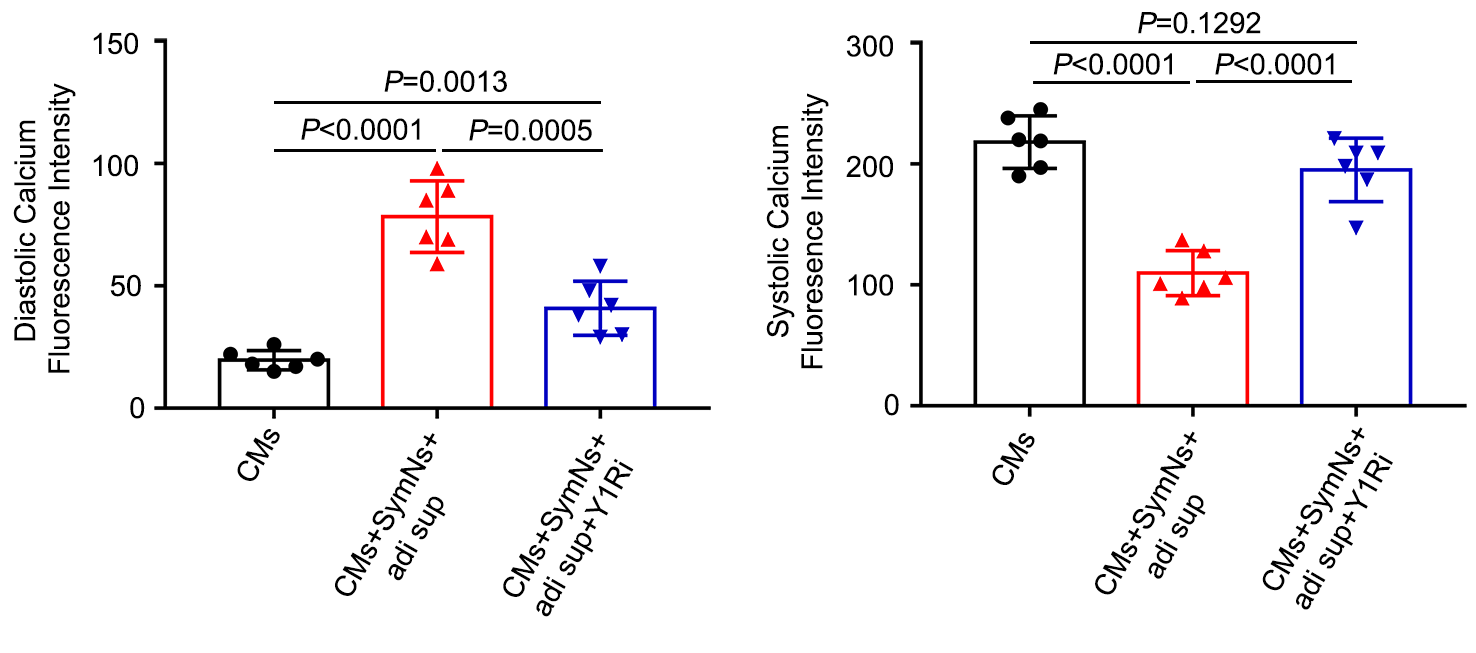

**Figure S8 The intracytoplasmic concentration of diastolic or systolic calcium of cardiomyocytes in triple co-culture system with or without Y1Ri.**

All of the error bars represent mean ± SEM.
